# Pre-challenge gut microbial signature predicts RhCMV/SIV vaccine efficacy in rhesus macaques

**DOI:** 10.1101/2024.02.27.582186

**Authors:** Hayden N. Brochu, Elise Smith, Sangmi Jeong, Michelle Carlson, Scott G. Hansen, Jennifer Tisoncik-Go, Lynn Law, Louis J. Picker, Michael Gale, Xinxia Peng

## Abstract

**Background:** RhCMV/SIV vaccines protect ∼59% of vaccinated rhesus macaques against repeated limiting-dose intra-rectal exposure with highly pathogenic SIVmac239M, but the exact mechanism responsible for the vaccine efficacy is not known. It is becoming evident that complex interactions exist between gut microbiota and the host immune system. Here we aimed to investigate if the rhesus gut microbiome impacts RhCMV/SIV vaccine-induced protection.

**Methods:** Three groups of 15 rhesus macaques naturally pre-exposed to RhCMV were vaccinated with RhCMV/SIV vaccines. Rectal swabs were collected longitudinally both before SIV challenge (after vaccination) and post challenge and were profiled using 16S rRNA based microbiome analysis.

**Results:** We identified ∼2,400 16S rRNA amplicon sequence variants (ASVs), representing potential bacterial species/strains. Global gut microbial profiles were strongly associated with each of the three vaccination groups, and all animals tended to maintain consistent profiles throughout the pre-challenge phase. Despite vaccination group differences, using newly developed compositional data analysis techniques we identified a common gut microbial signature predictive of vaccine protection outcome across the three vaccination groups. Part of this microbial signature persisted even after SIV challenge. We also observed a strong correlation between this microbial signature and an early signature derived from whole blood transcriptomes in the same animals.

**Conclusions:** Our findings indicate that changes in gut microbiomes are associated with RhCMV/SIV vaccine-induced protection and early host response to vaccination in rhesus macaques.

## INTRODUCTION

Live-attenuated rhesus cytomegalovirus-based SIV (RhCMV/SIV) vaccines provide sustained, durable protection against SIV challenge in ∼59% of vaccinated rhesus macaques [1,2], and it is unclear whether gut microbiota influence vaccine efficacy. In this study, we sought to interrogate gut microbiome profiles of RhCMV/SIV vaccinated rhesus macaques and determine if microbial RhCMV/SIV correlates of immunity exist.

The gut microbiome plays a significant role in host immunity and homeostasis, regulating response to infection and many autoimmune diseases [3], such as asthma [4,5] and Crohn’s disease [6]. The gut microbiome has also been shown to module immune repertoires and helper T cells in germ-free mice during early immune system development [7]. During acute HIV infection, gut barrier dysfunction is known to occur, resulting in increased microbial translocation in a condition known as ‘leaky gut’ [8]. This microbial translocation has been associated with immune response activation and inflammation during HIV infection [9] and was also shown to persist in some patients with poor immune response to antiretroviral therapy (ART) [10]. There is now evidence that gut microbiota are associated with vaccine-induced immunity in multiple species, including human and rhesus macaque [11]. It was recently shown using 16S rRNA sequencing that the gut microbiome is associated with protection outcome of rhesus macaques vaccinated with Adenovirus/protein simian immunodeficiency virus (SIV) vaccines [12,13]. A separate study revealed a strong gut microbiome response to administration of an HIV-1 DNA vaccine [14]. Whether or not the gut microbiome might influence RhCMV/SIV vaccine efficacy is still unknown.

16S rRNA gene sequencing provides the ability to rapidly profile microbiomes and determine these potential associations with host immunity. Amplicon sequence variants (ASVs) enable finely resolved profiling of community species/strains [15,16], but differential abundance analysis (DA) of this data is challenging due to its sparsity and the disagreement in results yielded by many available tools [17,18]. There is growing support for machine learning-based approaches that could replace or complement DA analysis of microbiome data [19,20]. Further, it is now critical to treat microbial count data as compositional using log-ratios [21], but common log-ratio techniques, such as centered log-ratios (CLRs), are prone to asymmetries in microbial data [22] and are collinear, complicating their statistical analysis [23,24]. There is also a suite of alternative techniques tailored for microbiome analysis [25–27], some of which explore the utility of phylogenetic log-ratios [28,29], but these complex log-ratios are challenging to interpret.

In this study, we analyzed the pre- and post-challenge gut microbiome profiles of RhCMV/SIV vaccinated rhesus macaques. We employed multiple compositional data analysis techniques available [30] and further developed log-ratio-based techniques to generate reference frames with properties similar to CLRs and avoid collinearity. We also developed a pairwise log-ratio-based feature reduction method and use this in combination with machine learning techniques. Using these methods, we analyzed 16S rRNA sequencing data generated from the gut microbiomes of RhCMV/SIV vaccinated rhesus macaques, revealing a pre-challenge gut microbial signature that predicts RhCMV/SIV protection outcome. We also used a common compositional DA analysis approach to corroborate these findings. We further show that this microbial signature significantly correlates with an independently identified protective signature in whole blood [31]. These findings show that the gut microbiome may be involved in RhCMV/SIV vaccine-induced immunity and warrant further research into the mechanisms underlying this relationship.

## RESULTS

### Rhesus gut microbiome profiles are diverse across different individuals but remain relatively stable within the same individuals during pre-challenge phase

We analyzed the gut microbiomes of rhesus macaques from three vaccination groups (groups O, S, and X) ([31] and **Figure 1a**). Each group had 15 animals and ∼55% of vaccinated animals were protected against repeated low dose SIV challenges (**Figure 1a**). Rectal swab samples for each animal were collected longitudinally after vaccination but before SIV challenge, with 5 time points for each animal in 3 week intervals (**Figure 1b**, ‘pre-challenge’ samples). Rectal samples were also continuously collected after SIV challenge for 27 animals (23 protected, 4 not protected), spanning 28 to 252 days after the animals became infected (**Figure 1b**, ‘post-challenge’ samples). While only 4 of the 22 not protected animals were sampled post-challenge, each vaccination group was represented among the selected animals (1 O, 1 S, and 2 X group animals). We profiled the microbiome compositions in all rectal samples using 16S rRNA gene V4 region-based sequencing analysis and obtained ∼52.5 million paired-end reads across 220 pre- and 129 post-challenge samples (∼150,000 reads per sample). Among these samples, we detected ∼2,400 unique ASVs representing communities of potential bacterial species/strains. A total of 43.5M of 52.5 M reads (∼82.9%) were merged and assigned to these ASVs, resulting in ∼125,000 ASV read counts per sample. As expected, the most prevalent phyla in the samples were Spirochaetes, Firmicutes, and Bacteroidetes, regardless of vaccination group, protection outcome, or challenge phase (**Figure 1c**). In some animals, we observed a high proportion of Campilobacterota that was also irrespective of study covariates, where Campilobacterota constituted >50% of all bacteria detected, although this was uncommon (26 of 347 samples, 7.5%) (**Figure 1c**).

**Figure 1.**
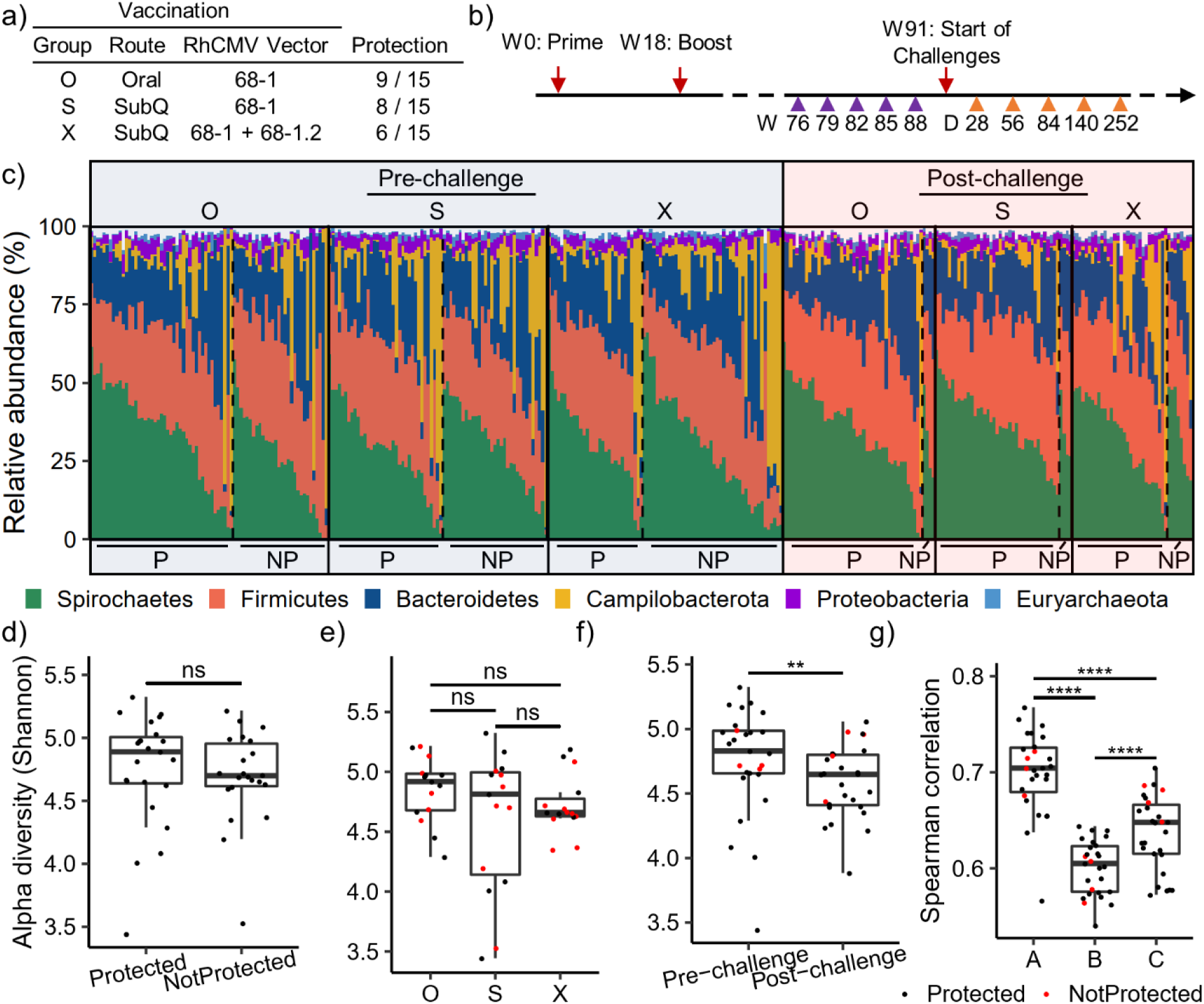
Overview of rhesus gut microbiomes from RhCMV/SIV vaccine study. **a)** Vaccination groups showing vaccine routes (Oral or SubQ = subcutaneous), RhCMV vector(s), and the protection outcomes of animals. **b)** Time course of the study, showing the weeks (W) that pre-challenge samples were collected, relative to vaccination prime and boost. Post-challenge samples are shown as days (D) after the animal was infected. **c)** Relative abundances of phyla in each individual sample, organized by challenge phase (pre or post), vaccination group (O, S, X) and protection outcome (P = protected, NP = not protected). Vertical solid lines separate vaccination groups and vertical dotted lines separate P/NP animals within groups. Samples are further organized by their animal label (i.e. samples from the same animal are adjacent stacked bars). Only phyla with >= 1% average relative abundance are shown. **d-f)** Average alpha diversities computed using the Shannon index are shown for animals stratified over protection outcome **(d)**, vaccination group **(e)**, and challenge phase **(f)**. **g)** Average spearman correlations are computed for each animal using pre-challenge samples (A), post-challenge samples (C), and strictly between pre- and post-challenge samples (B). Where applicable, animals are colored if by protected outcome (P = black, NP = red) (e-g). In **(f-g)**, animals are limited to those with both pre- and post-challenge samples. Statistical significance was determined using two-sided unpaired (**d**, **e**, and **g**) and one-sided paired (**f**) t-tests. ****p < 0.001, ***p < 0.01, **p < 0.05, *p < 0.1, ns = not significant.

We first examined potential changes in alpha diversities across these microbiome profiles. While we did not observe a significant difference in pre-challenge alpha diversity based on protection outcome or vaccination group (**Figure 1d and 1e**), there was a significant decrease after SIV challenge (**Figure 1f**, p < 0.05, one-sided paired t-test), in line with previous findings of SIV-induced gut dysbiosis [32]. We then compared the similarities of the microbiome profiles from the same animals before and after SIV challenge. We also observed that average correlations among each animal’s pre-challenge samples (i.e. between an animal’s different time points) were significantly higher than those of post-challenge samples (**Figure 1g**, p < 0.001, two-sided paired t-test), suggesting that the gut microbiome was more dynamic after SIV challenge. The correlations between pre- and post-challenge samples from the same animal were even lower (**Figure 1g**). This shows that SIV challenges significantly impacted the animal gut microbiomes. Since most (23 of 27) of the post-challenge samples were from protected animals, the changes in microbiome compositions induced by SIV challenge were still not fully restored even long after the animals cleared infection.

Overall, the microbiome profiles from the same animals collected at different time points were highly similar to each other (average spearman correlation coefficients around 0.6-0.7), suggesting the overall microbiome profiles within each animal tended to be stable during the whole vaccine study. When all samples were clustered together based on pairwise correlations, samples from the same animal, both pre- and post-challenge, in general tended to be grouped close to each other (**Figure S1**). This further shows the overall microbiome profiles within each individual remained mostly stable, relative to the large between-animal differences. This observed robustness of microbiome profiles within animals motivated us to aggregate count data across animal time points within each phase of the study, resulting in one aggregated pre-challenge read count and one aggregated post-challenge read count for each animal (Methods). We further filtered this aggregated dataset by requiring 5 counts in 50% of animals, yielding 1,434 ASVs for downstream analysis. This process drastically reduced data sparsity and yielded an aggregated microbiome count matrix with only 6.2% zero counts, while unaggregated microbiomes had 27.4% zero counts (**Supplementary Data**). This low prevalence of zero counts yielded a dataset better suited for downstream compositional analysis.

### Microbiota reference frames reveal a strong vaccination group effect

We next investigated if these pre-challenge microbial profiles distinguished vaccination groups. Since animals from different vaccination groups were vaccinated using different administration routes and/or different vaccine vectors (**Figure 1a**), we sought to determine if these group differences might alter gut microbiome compositions as well. We first used a phylogeny-based approach to construct balances (i.e. isometric log-ratios) of ASVs [28], where each balance was a node in the phylogenetic tree and the log-ratio numerator and denominator were composed of descendants on either side of the node. Since pairwise balances at the bottom of the tree tend to be more variable, as previously observed [28], we selected the most variable pairwise balances from the bottom of the tree (Methods) and used them for principal components analysis (PCA) (**Figure S2a-c**). Interestingly, we found significant separation of vaccination groups (p < 0.0001, PERMANOVA) using this approach (**Figure S2c**), suggesting that RhCMV/SIV vaccine administration route and/or vector(s) differentially altered rhesus gut microbiome compositions.

This strong signal motivated us to determine if we could identify more interpretable log-ratios that discriminate microbiomes by their vaccination groups. We began by identifying a set of ASVs to use as a reference frame for representation of the remainder of ASVs in compositional space to standardize ASV abundances across all samples (Methods and **Supplementary Methods**). CLRs use all ASVs as a reference frame which has zero variance by definition, and we used a Markov chain Monte Carlo (MCMC) procedure to approximate CLRs only using a parsimonious set of ubiquitously detected ASVs. In our MCMC procedure, we repeatedly sampled highly abundant ASVs, with the objective of minimizing the variance of the resulting reference frame geometric mean. Each MCMC iteration consisted of first sampling a set of ASVs and then adding or subtracting ASVs to reduce the variance of the geometric mean of the set until a local minimum was reached. Using 5 MCMC experiments each with 500 iterations, we identified multiple stable reference frames as candidates for standardization (**Figure S3a-b**). On average, MCMC-generated reference frames were composed of only 90 ASVs and the number of transitions in MCMC iterations depended on the size of the initial ASV set randomly selected (**Figure S3c-e**). Using loess regression we modeled the variances of the 2,500 reference frames produced by these MCMC experiments as a function of their size and identified the optimal parsimonious and low-variance reference frame from each experiment (**Figure S3f**). These reference frames contained many of the same ASVs (22), though reference sets also had unique ASVs and ASVs common to only 3 of 4 of the sets (ranging from 0 to 16 ASVs), suggesting possible degeneracy of viable reference ASVs (**Figure S3g**). These reference sets also produced standardized ASV counts similar to each other and to the CLR, confirming that these reference sets indeed have similarly low variance (**Figure S3h**). A final reference frame of 69 ASVs was selected with optimal residual value from our regression (**Figure S3f**), composed primarily of bacteria from the *Lachnospiraceae* and *Ruminococcaceae* families, which are essential commensals in mammalian gut microbiomes that metabolize plants [33] (**Supplementary Data**).

Within each sample we standardized the remaining 1,365 ASVs as log-ratios with respect to the geometric mean of the reference frame, yielding standardized values with normal properties highly similar to those of the centered log-ratio (**Figure 2a**). Strikingly, ∼55% of the animal-to-animal variation was explained using PCA with only the top five variable ASVs (**Figure 2b**), which also significantly segregated animals by vaccination group (**Figure 2c**, PERMANOVA, p < 0.0001). Four of the top five variable ASVs were taxonomically assigned to the classes Clostridia (ASV53, ASV84), Sphingobacteriia (ASV64) and Bacteroidia (ASV263), while the fifth was *Methanomassiliicoccus luminyensis* (ASV277) (**Supplementary Data**), a methanogenic archaeon found in the human gut [34]. Interestingly, ASV277 had stronger detection in group X (-0.536) compared to groups O and S, with an average log-standardized value of -3.7 (∼4.6 log2 fold difference). *Kineothrix alysoides* (ASV53) was also much higher in group X, with an average log-standardized difference of 2 relative to groups O and S. We also found a mutually exclusive detection of Bacteroidetes species depending on the animal’s vaccination route, strongly detecting ASV263 only in group O and ASV64 exclusively in groups S and X. Lastly, a second species from Clostridia (ASV84) had a range of detection based on vaccination group, most prominent in group O followed by group X then S. These results of the overall differences in vaccination groups corroborate the phylogenetic-based findings (**Figure S2**) and also elucidate the differences in individual gut microbiome features that existed between the three vaccination groups pre-challenge.

**Figure 2.**
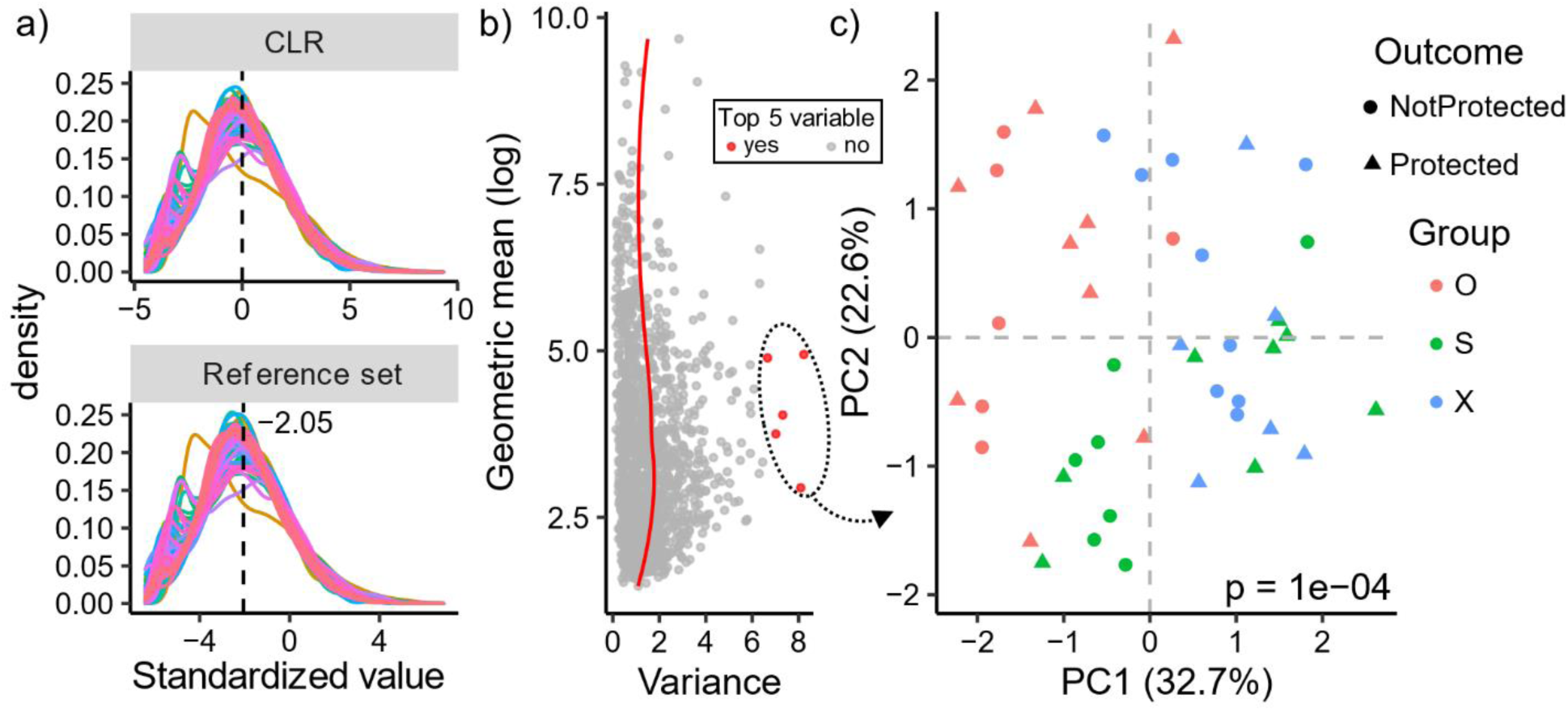
Highly variable, standardized ASVs distinguish rhesus gut microbiomes based on vaccination groups. **a)** Standardization of ASVs using a reference frame of 69 ASVs yields similar normal properties to the centered log-ratio (CLR). Standardization with the reference set does not guarantee centering at zero, and the mean centering is shown. **b)** Scatter plot of mean-variance trend of reference standardized ASVs with loess regression curve shown in red. The top five variable ASVs are shown in red and used for PCA in **(c)**, where animals are colored by vaccination group and have shapes based on protection outcome. Statistical significance was determined using PERMANOVA.

### Pre-challenge gut microbiomes predict animal vaccination groups

This clear vaccination group effect prompted us to examine the gut microbiomes more closely to identify a more comprehensive list of vaccination group associated ASVs. We used multiple strategies to build random forest classifiers designed to separate one vaccination group from the other two (Methods). Briefly, three types of ASV standardized values were used in this analysis: 1) centered log-ratio (CLR) standardized ASVs, 2) reference log-standardized ASVs, and 3) phylogenetic balances. We also developed and applied a greedy pairwise log-ratio strategy (Pairbal) to dimensionally reduce the CLR and reference log-standardized values prior to using the random forest classifier (Methods). Model predictions were compared between experiments using this dimension reduction approach, a naïve dimension reduction approach using only the most abundant ASVs, and an experiment not using dimension reduction at all (Methods). These classifiers had strong predictive power regardless of the vaccination group comparison, with average area under the curves (AUCs) of 0.924-0.991 when predicting animals in the pre-challenge phase (**Table 1**, **Figure S4**). Interestingly, these classifiers trained with pre-challenge samples also performed well when predicting vaccination groups of animals using post-challenge samples, with mean AUCs ranging from 0.726 to 0.903 (**Figure S4**). Phylogenetic balances as input to classifiers had the strongest average area under the curve (AUC) when comparing X against O and S; meanwhile reference log-standardization performed best with distinguishing O from S and X and CLRs best distinguished S from O and X (**Table 1**). We further found that all CLR and custom reference frame models using only Pairbal selected ASVs as input had significantly increased AUC relative to models using either all ASVs or the most abundant ASVs (**Table 1**, **Figure S4**).

**Table 1.** 15-fold cross-validation (CV) using random forest classifiers to predict animal vaccination groups. Random forest classifiers were constructed using pre-challenge samples and the vaccine group of animals left out were predicted using either their pre- or post-challenge samples. Classifiers were constructed in a binary fashion, comparing one vaccination group against the other two, yielding three comparisons total. Different microbial features were used as input to the classifiers. Centered log-ratio (CLR) standardized and reference log-standardized ASVs were used as input to the classifiers either using all ASVs, most abundant ASVs, or only those selected by Pairbal. The number of most abundant ASVs chosen matched the number selected by Pairbal. Phylogenetic balances constructed using the PhILR tools are also shown for comparison. In all cases, the area under the curve (AUC) was computed by taking the average of the five 15-fold CV AUCs.

| Study phase | Comparison | CLR |  |  | Reference |  |  | PhILR |
| --- | --- | --- | --- | --- | --- | --- | --- | --- |
|  |  | all | abundant | Pairbal | all | abundant | Pairbal |  |
| pre-challenge | X vs O,S | 0.954 | 0.897 | 0.976 | 0.949 | 0.911 | 0.976 | **0.988** |
|  | S vs O,X | 0.928 | 0.857 | **0.955** | 0.925 | 0.943 | 0.951 | 0.940 |
|  | O vs S,X | 0.964 | 0.920 | 0.975 | 0.970 | 0.941 | **0.979** | 0.967 |
| post-challenge | X vs O,S | 0.796 | 0.565 | 0.853 | 0.807 | 0.772 | 0.842 | **0.903** |
|  | S vs O,X | 0.769 | 0.757 | **0.830** | 0.785 | 0.796 | 0.827 | 0.724 |
|  | O vs S,X | 0.832 | 0.723 | **0.882** | 0.841 | 0.738 | 0.878 | 0.802 |

Next, we more closely analyzed the ASVs identified by Pairbal cross-validation (CV) experiments that we used to examine differences between the vaccination groups. Each of the 5 CV experiments were 15-fold (i.e. leaving three animals out each time), yielding a total of 75 prediction experiments, where an ASV might be included in the Pairbal selection. We found that Pairbal identification frequency largely followed a bimodal distribution (**Figure 3a**). The lower tail of the distribution represented ASVs infrequently used by predictive models, while those appearing in the upper tail were more commonly used. Since Pairbal enhanced predictive model performance (**Figure S4**), we took a consensus CV approach, observing 284 ASVs identified in all Pairbal selections in at least one of the group comparisons (consensus CV ASVs). These ASVs included the top five variable ASVs previously shown to segregate animals by vaccination group (**Figure 2b**, **Supplementary Data**). Interestingly, consensus CV ASVs significantly segregated animals by vaccination groups using both pre- and post-challenge samples (**Figure 3b**, PERMANOVA, p < 0.0001), indicating that pre-challenge vaccination group differences persisted after challenge. We then compared the vaccination group comparisons where consensus CV ASVs were identified in all 75 Pairbal selections, finding that many of the ASVs were identified in at least two of the three comparisons (**Figure 3c**, 135 of 284, 47.5%). The comparison between X with O and S yielded 74 unique ASVs, the largest number of unique ASVs among the three vaccination group comparisons, suggesting that animals in group X may have had gut microbiomes further diverged from the other two groups in general. Taxonomic classifications of the consensus CV ASVs revealed that most were assigned to the Clostrida class (**Figure 3d**, 148 of 284, 52.1%). Of ASVs from Clostridia, 60 unique genera were identified with many ASVs from *Ruminococcus* (17 ASVs), *Clostridium sensu stricto* (8 ASVs), and *Lachnospiracea incertae sedis* (7 ASVs) (**Figure 3e**). Other common classes were Bacterodia (31 ASVs), Deltaproteobacteria (20 ASVs), Alphaproteobacteria (18 ASVs), and Mollicutes (16 ASVs) (**Figure 3d**).

**Figure 3.**
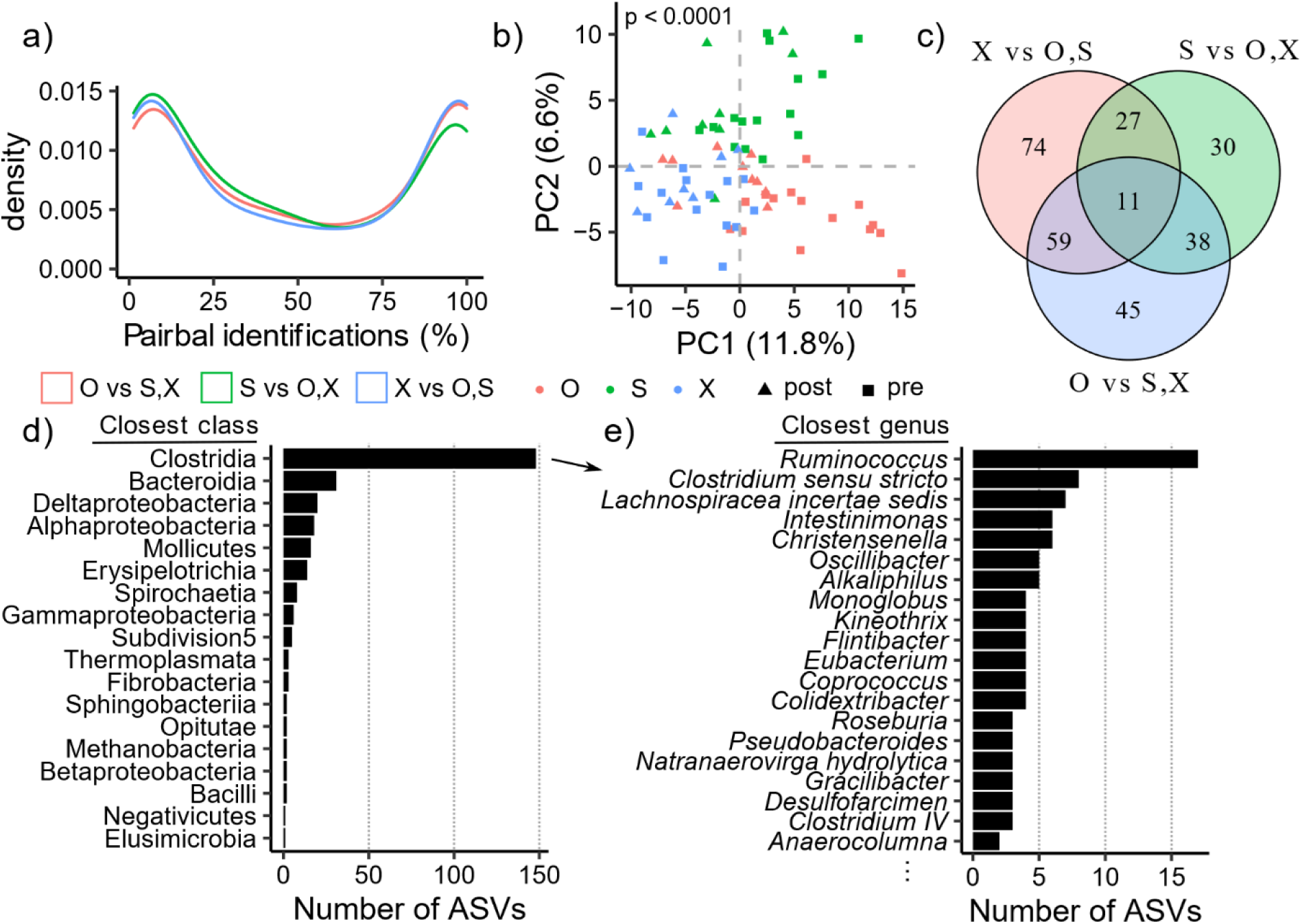
Detection of a vaccination group associated microbial signature. Vaccination groups are grouped together for binary predictions, e.g. in O vs S,X the S and X groups are combined. Random forest predictions for pre- and post-challenge samples were generated using 5 iterations of 15-fold cross-validation (CV) experiments using only pre-challenge samples. Before training each random forest model, Pairbal was used to dimensionally reduce the training data. **a)** Distribution of ASVs based on the proportion of CV predictions (i.e. predictive models) where they were identified by Pairbal, where lines are colored based on the comparison. **b)** Principal components analysis using the 284 ASVs identified in all predictions, with statistical significance of vaccination group effect assessed using PERMANOVA, p = 0.0001. **c)** Venn diagram showing the ASVs identified in each comparison. **d)** Bar plot showing the number of ASVs assigned to each class. **e)** Bar plot showing the number of ASVs assigned to the top 20 most frequent genera detected within Clostridia.

While we observed a consistently strong vaccination group effect in our initial ASV variation analysis (**Figure 2c**) and via random forest analyses utilizing various microbial feature types (**Table 1**, **Figure S4**, **Figure 3**), we sought to confirm these findings independently using ANCOM-BC, which is a more recently developed differential abundance (DA) analysis method [35]. Using the same vaccination group comparison approach (O vs. S and X, etc.), we found that the group associated consensus CV ASVs from Pairbal had a statistically significant propensity to have higher absolute log2 fold-changes (L2FCs) and lower FDR-adjusted p-values identified by ANCOM-BC compared to all other ASVs (**Figure S5a-d**, **Supplementary Data**; Wilcoxon rank-sum tests, both p < 0.001). In fact, a total of 41 significantly DA ASVs were found across the three comparisons by ANCOM-BC, most of which were among the consensus CV ASVs (**Figure S5e**, 38, 92.7%). Visualization of the DA L2FCs confirmed that Pairbal identified ASVs that exhibited distinct abundance patterns across vaccination groups, which loosely formed five groups broadly annotated as high, medium, and low depending on L2FCs observed across the three vaccination groups (**Figure S6**). For example, 8 ASVs (group 4, **Figure S6**) were enriched in group S relative to groups O and X, comprising three strains of *Ruminococcus* (**Supplementary Data**). Another group of 111 ASVs (group 3, **Figure S6**) were mostly from Clostridia (59 ASVs) and enriched in group X relative to groups O and S (**Supplementary Data**). These results further show the gut microbiome differences between vaccination groups. Since these rectal samples were collected at least 58 weeks after the last boost, these results also suggest that the potential impact of vaccination route and/or vectors on the compositions of gut flora could persist long-term.

### Pre-challenge gut microbial protection signature predicts vaccine efficacy

Given that we observed a significant pre-challenge vaccination group associated microbial signature, we hypothesized that there might also be pre-challenge protection associated microbial signatures across the three vaccination groups. We applied the same strategy as with our vaccination group comparisons, using 1) random forest classifiers with CLRs, 2) reference log-standardized values, and 3) phylogenetic balances as input. Similarly, we also applied our feature reduction strategy, Pairbal. We again used 5 iterations of 15-fold CV experiments (Methods). Using these different microbial feature inputs we found that using Pairbal, in conjunction with either reference log-standardization or CLRs, resulted in the highest AUC (∼0.77 on average) when the pre-challenge samples were used for predicting vaccine protection outcome (**Figure S7**). These two approaches performed similarly to phylogenetic balances when the post-challenge samples were used for prediction (∼0.73 AUC) (**Figure S7**). Further, Pairbal-based feature reduction greatly improved the performance of predicting protection outcomes using pre-challenge samples, with significantly higher AUCs than predictive models using either all ASVs or the most abundant ASVs (**Figure S7**, one-sided paired t-tests, all p < 0.05).

We further examined the prediction results with Pairbal used in conjunction with reference log-standardization, where we observed robust prediction of protection outcome using either pre-challenge samples (AUC = 0.776) or post-challenge samples (AUC = 0.731) (**Figure 4a-b**). Similar to the analysis of vaccination group-associated microbiota (**Figure 3a**), we observed that the frequency of ASVs identified by Pairbal predictive models followed a bimodal distribution, albeit with an upper tail of lower density (**Figure 4c**). Using the 47 ASVs identified in >95% of predictive models (consensus CV ASVs), we found that there was a significant separation of animals based on protection outcome in the pre-challenge phase (**Figure 4d**, PERMANOVA, p < 0.0001). Nearly half of the consensus CV ASVs were from Clostridia (20 of 47, 42.6%) (**Figure 4e**), harboring a variety of genera (17 total), with *Clostridium sensu stricto* (2 ASVs), *Intestinimonas* (2 ASVs), and *Ruminococcus* (2 ASVs) chief among them (**Figure 4f**). Overall, 29 of 47 (61.7%) of the protection associated consensus CV ASVs were not among the 284 vaccination group associated consensus CV ASVs (**Figure 4e**). Interestingly, most protection associated Clostridia ASVs were not identified as vaccination group associated (**Figure 4e**, 14 of 20, 70%).

**Figure 4.**
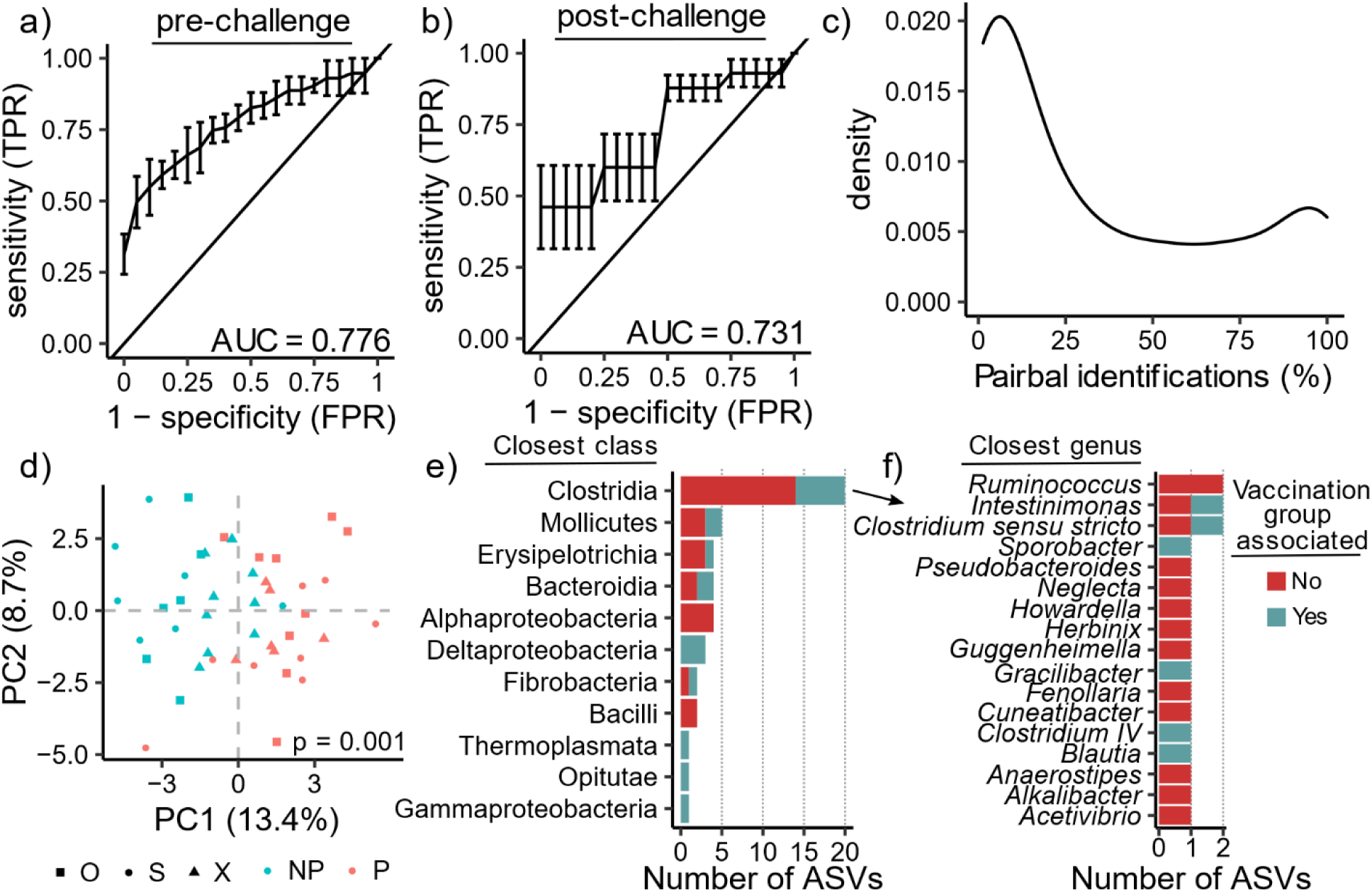
Detection of a protection associated microbial signature. Random forest predictions for pre- and post-challenge samples were generated using 5 iterations of 15-fold cross-validation (CV) experiments using only pre-challenge samples. Before training each random forest model, a microbial reference frame was determined Pairbal was used to dimensionally reduce the training data (reference frame standardized ASV counts). Post-challenge samples were only predicted using models trained with pre-challenge samples from different animals. **a-b)** Receiver operating characteristic (ROC) curves with area under the curve (AUC) shown for prediction of pre-challenge **(a)** and post-challenge **(b)** samples. **c)** Distribution of ASVs based on the proportion of CV predictions (i.e. predictive models) where they were identified by Pairbal. d) Principal components analysis using the 47 ASVs identified in at least 95% of predictions. Significance was assessed using PERMANOVA, p = 0.001. **e)** Bar plot showing the number of ASVs assigned to each class. **f)** Bar plot showing the number of ASVs assigned to the genera detected within Clostridia. In **(e-f)**, bars are colored based on whether the ASV was vaccination group-associated (blue) or not (red).

Similar to our vaccination group analysis, we performed DA analysis using ANCOM-BC to independently evaluate these Pairbal random forest findings. The 47 ASVs identified by Pairbal in >95% of CV predictive models had a statistical significant propensity to be among ASVs with higher absolute L2FCs and lower raw p-values as determined by ANCOM-BC (**Figure S8a-d**, **Supplementary Data**; Wilcoxon rank-sum tests, both p < 0.01). While ANCOM-BC did not identify any significantly DA ASVs (i.e. no ASVs with both absolute L2FC > 1 and FDR < 0.05), there was a propensity for Pairbal to detect ASVs with raw p-values < 0.05 (11 of 28, 39.2%) and absolute L2FCs > 1 (25 of 64, 39.1%) (**Figure S8e**). Visualization of ASV L2FCs revealed that ASVs not detected by Pairbal that had a raw p-value < 0.05 formed a second small cluster of 17 ASVs, implying that some ASVs below the 95% Pairbal identification threshold used might also be associated with protection (**Figure S9**). We found that these ASVs were commonly from Clostridia regardless of whether they were enriched (8 of 12, 66.7%) or depleted (4 of 5, 80%) in protected animals, similar to the ASVs found by Pairbal (**Figure 4e**). Many of these DA Clostridia ASVs were detected by Pairbal in >50% of Pairbal identifications (5 of 12, 41.7%). Together, these results show that a common pre-challenge protective signature existed across the three vaccination groups despite the clear between-group differences we observed.

### Features of pre-challenge protective microbial signature

We next examined how these 47 ASVs varied across the 45 animals during the pre-challenge phase. Visually we observed similar trends in the microbial relative abundances across the different vaccination groups with some animal-to-animal variation (**Figure S10**), which we expected based on our initial analysis of animal gut microbiome profiles (**Figure S1**). Most of these ASVs (29 of 47, 61.7%) were relatively more abundant among protected animals (**Table S1**, **Supplementary Data**). Furthermore, 26 ASVs were found to have consistent enrichment (19 ASVs) and depletion (7 ASVs) across all three vaccination groups. We also examined the phylogenetic relationship of these ASVs and visually observed groups of closely related ASVs that were collectively enriched or depleted among protected animals (**Figure S11**). For example, we identified 6 neighboring ASVs in the phylogeny from Clostridia that were depleted among protected animals, while we also observed 6 neighboring ASVs from Proteobacteria that were enriched among protected animals (**Figure S11**).

We then sought to determine if SIV challenge had any effect on the relative abundances of these protection associated ASVs. Since the majority of our post-challenge samples were from protected animals (23 of 27, 85%), we elected to exclude all four not protected animals from this analysis. Before beginning our analysis, we confirmed that our reference set of 69 ASVs similarly centered post-challenge samples, finding no difference in the overall distributions between pre- and post-challenge samples of those protected animals (**Figure 5a**). Three Clostridia ASVs depleted among protected animals pre-challenge significantly increased after SIV challenge (**Figure 5b**, one-sided paired t-test, FDR < 0.05). Relaxing the significance threshold to FDR < 0.1, four additional depleted Clostridia ASVs were observed to increase after SIV challenge and a diverse group of five enriched ASVs decreased (**Figure 5b**). This limited perturbation of protection associated ASVs was consistent with our earlier finding that the predictors learned from pre-challenge samples were also able to predict protection status using post-challenge samples as input (**Figure 4d** and **Figure S7**). Collectively, these results indicate that while some pre-challenge protection associated ASVs were altered by SIV challenge, most persisted in the same animals long after SIV challenge.

**Figure 5.**
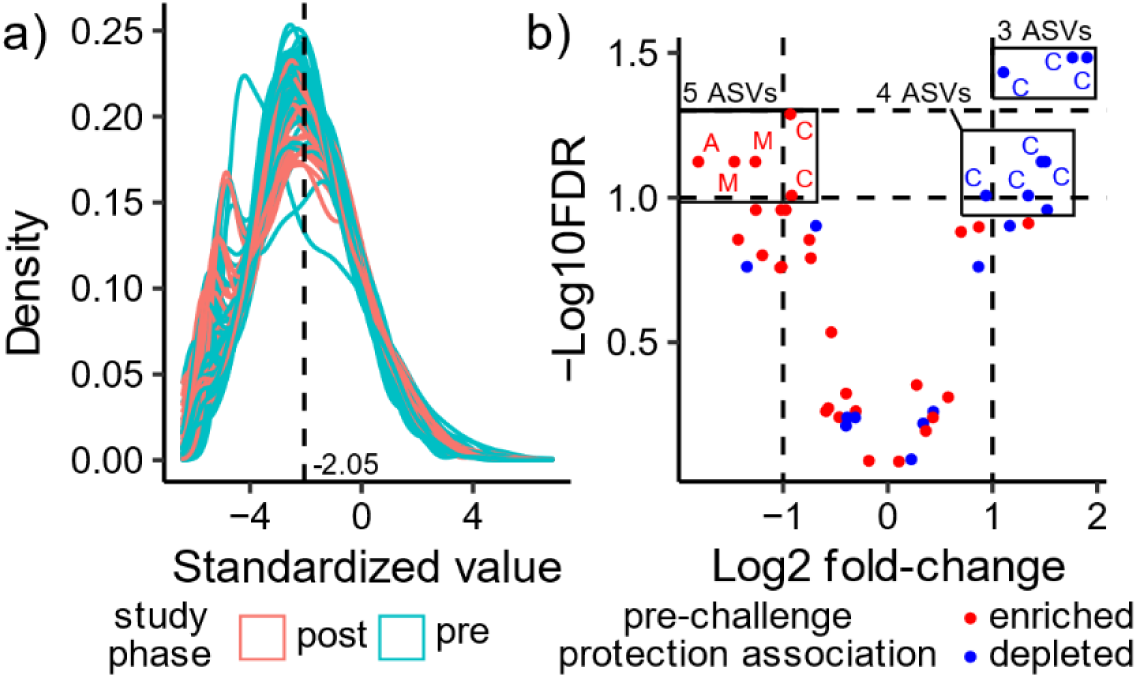
Perturbations of protection associated ASVs after SIV challenge. **a)** Standardization of ASVs using a reference frame of 69 ASVs, with pre-challenge (pre) and post-challenge (post) samples colored blue and red, respectively. **b)** Volcano plot of protection associated ASVs post-vs. pre-challenge. Log2 fold-change (L2FC) and -log10 FDR are shown for all 47 ASVs. ASVs are colored based on whether they are enriched (red) or depleted (blue) in protected animals pre-challenge. The two horizontal dashed lines indicate FDR thresholds of 0.05 (top) and 0.1 (bottom), and the two vertical dashed lines indicate an absolute L2FC > 1. ASVs with FDR < 0.1 are labeled by the first letter of their taxonomic class: A=Alphaproteobacteria, C=Clostridia, M=Mollicutes. Statistical analysis was performed using one-sided paired t-tests with FDR control.

### Protection signatures from gut microbiomes and host whole blood are significantly correlated

Previously, using mRNA-seqanalysis of whole blood samples from the same animals (vaccination groups O and S) we observed a sustained, coordinated immune response regulated by Interluekin-15 during RhCMV/SIV vaccination that mediates protection outcome [31]. The whole blood samples used in that analysis were collected during vaccination (W0 and W18), well before the rectal swabs were collected for the analysis of gut microbiomes here (**Figure 1b**). We thus hypothesized that the microbial protective signature we observed in this study might be ‘imprinted’ by those earlier protection signatures identified in whole blood. We thus investigated the correlations between the abundance of protection associated ASVs with the vaccination vs. baseline expression changes of differential differentially expressed (DDE) genes. DDE genes were defined as genes with significantly different (absolute L2FC > 1.5 and FDR < 0.05) baseline-subtracted gene expression between protected and not protected animals. Thus, positive DDE fold changes indicate genes with larger changes in expression relative to baseline in protected versus not protected animals. We assessed the significance of these correlations using permutation-based analyses, where we permuted genes, ASVs, and the animal labels separately. For a configuration of DDE genes with protection-associated ASVs to be considered significant, all three permutation-based p-values were required to be less than 0.05 (Methods). When we examined these DDE genes across all animals together, we found a significant overall correlation at W0D3, with empirical p-values of 0 from all three permutation analyses (**Figure S12**). We also found a significant, yet weaker correlation at W18D1 (gene, ASV empirical p-values < 0.001, animal empirical p-value < 0.05).

We then visualized the correlations between the DDE genes and ASVs at W0D3 where we observed the most significant correlations (**Figure 6**). An expected pattern emerged where DDE genes with negative fold changes were positively correlated with ASVs that were depleted in protected animals. Similarly such genes were negatively correlated with ASVs that were enriched in protected animals. This same pattern was found for genes with positive fold changes as they were positively and negatively correlated with ASVs that were more and less abundant in protected animals, respectively. Upon closer inspection of the dendrogram, we found two clusters of ASVs following this pattern: 14 of the 29 ASVs with positive associations (left cluster, bolded red ASV labels), and 10 of the 18 ASVs with negative associations (right cluster, bolded blue ASV labels) (**Figure 6**). Most of the ASVs in each cluster were uniquely protection associated without group association (10 of the 14 positively associated ASVs, 6 of the 10 negatively associated ASVs). In general, many of these ASVs were assigned to the Clostridia class (13 total), including 5 enriched and 8 depleted ASVs in protected animals. This community of Clostridia was also quite diverse, representing 12 different genera. Of these ASVs, ASV1489 (*Blautia obeum*, depleted in protected animals) had the strongest correlations with DDE gene expressions (median correlation coefficients of 0.345 and -0.298 with DDE genes of negative and positive fold changes, respectively). Together, these findings indicate that early vaccine-induced host response might have induced the protection-associated microbial signature we detected in pre-challenge samples collected long after vaccination.

**Figure 6.**
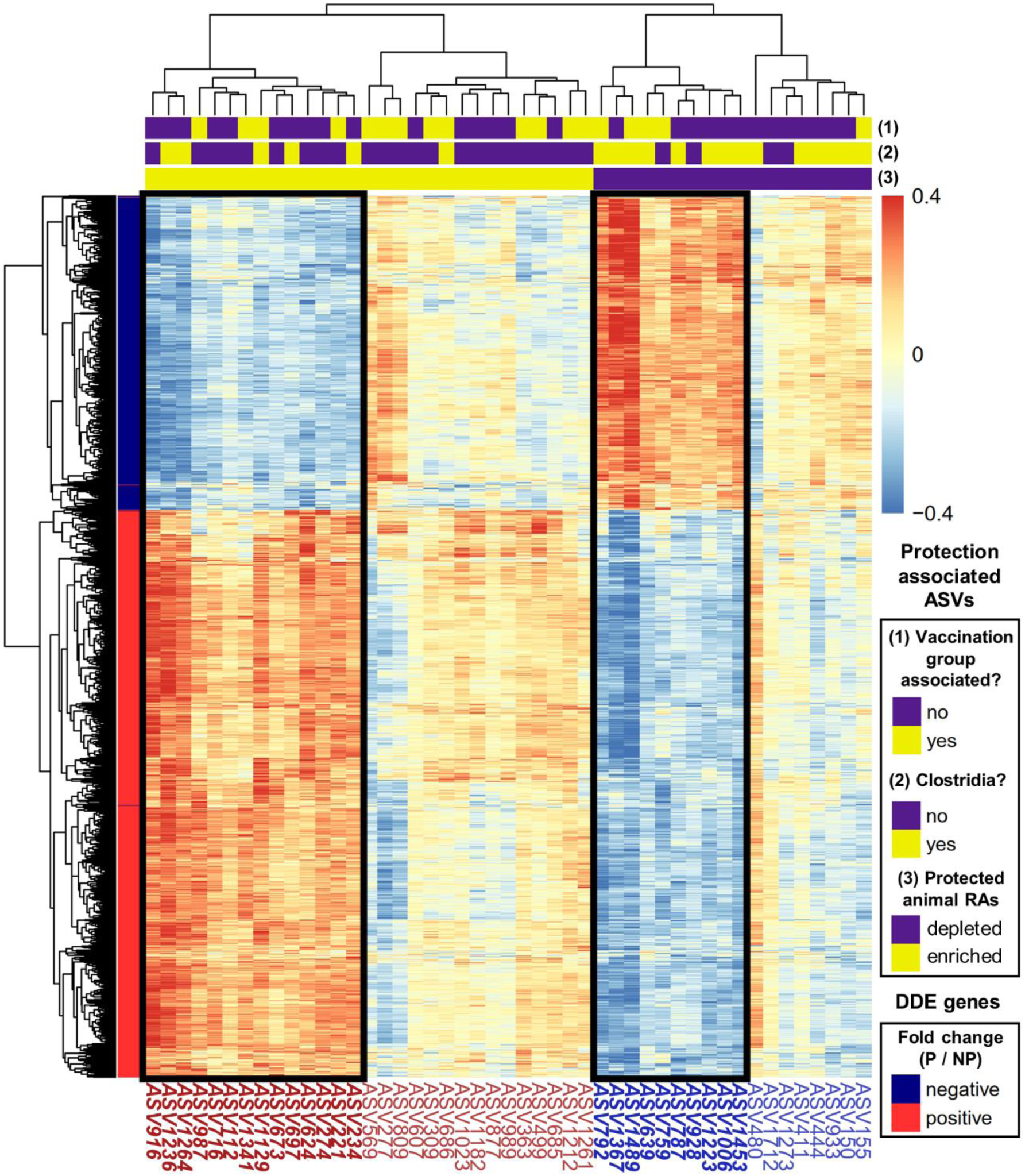
Correlations between protection associated ASVs and host DDE genes at W0D3. ASVs are clustered separately whether they are enriched (red labels, yellow in bottom bar above plot) or depleted (blue labels, purple in bottom bar above plot) in protected animals. ASVs are further colored based on whether they were taxonomically classified in the Clostridia class and whether they were associated with vaccination group (middle and top bars above plot, respectively: yes = yellow, no = purple). Similarly, DDE genes are clustered separately whether they have a positive (red) or negative (blue) fold change, respectively representing larger or smaller changes from baseline in protected (P) animals relative to non-protected (NP) animals. These are labeled with colored indicators to the right of the dendrogram. ASVs are labeled below the heatmap. Correlations are saturated such that correlations only range between +/- 0.4. Black boxes indicate clusters of ASVs with correlation patterns matching those that were expected given the configuration of DDE genes and protection associated ASVs with associated ASV labels below the plot bold and italicized. For example, DDE genes with positive fold changes were expected to positively correlate with protection associated ASVs with higher abundances in protected animals.

## DISCUSSION

### Association of gut microbiomes with RhCMV/SIV vaccine protection outcome in rhesus macaques

To our knowledge, this is the first study to investigate the association of gut microbiomes with RhCMV/SIV vaccine protection outcome in rhesus macaques. Using novel compositional data analysis techniques and a machine learning approach, we separately identified gut microbial signatures that were predictive of vaccination groups and vaccine protection outcomes. We further showed that these vaccine protection associated microbial features were significantly correlated with early vaccine-induced host immune signatures in whole blood from the same animals.

Here we presented three complementary pieces of evidence that the gut microbiome was likely involved in RhCMV/SIV vaccine-induced protection. First, we showed that gut microbiome profiles were significantly different across the three vaccination groups. Two of three groups were administered with the same 68-1 RhCMV/SIV vaccine, and only differed by routes of administration, i.e. subcutaneous vs. oral routes. The third group was vaccinated subcutaneously with a combined vaccine including the same 68-1 RhCMV/SIV vaccine and a variant 68-1.2 RhCMV/SIV vaccine with the same SIV inserts. All animals were naturally CMV exposed before vaccination and the samples for gut microbiome analysis were collected after vaccination but before animals were SIV challenged. These different RhCMV/SIV vaccines are the most plausible driver for the significant differences in gut microbiome profiles. Second, we identified a gut microbial signature from these pre-challenge samples that predicts vaccine protection outcome when animals were SIV challenged later, and this microbial signature was shared across the three vaccination groups. This is direct evidence that the changes in gut microbiomes are associated with RhCMV/SIV vaccine protection outcome. Interestingly, this predictive gut microbial signature was also relatively robust. The ASV abundances of ∼74% (35 of 47) of the features remained largely unchanged even after SIV challenge. The same signature was still predictive of vaccine protection outcome even when post-challenge gut microbiome profiles were used as input for prediction. However, we did observe perturbation of some protection associated microbes and an overall reduction in microbial composition and diversity after challenge, possibly suggestive of gut dysbiosis. This condition, commonly due to gut mucosal damage and thus referred to as ‘leaky gut’ [8], manifests in chronically infected HIV patients [9]. In SIV infected rhesus macaques, it was shown that gut dysbiosis occurs but can be partially reverted using combination antiretroviral therapy (c-ART) [36]. Third, we show that the vaccine protection associated microbial features were significantly correlated with the early vaccination-induced immune signatures identified in whole blood of the same animals. Importantly, the protection associated microbial signature was detected in pre-challenge samples that were collected long after vaccination. These findings show that early vaccination-induced host response might have been responsible for the long-lasting changes in gut microbiomes, wholly or in part.

It is unclear how the identified microbial features might mechanistically influence RhCMV/SIV vaccine protection. These signatures were largely comprised of bacteria from the Clostridia class. Interestingly, some bacterial strains from Clostridia (from clusters XIVa, IV and XVIII) can induce Treg cells in the murine gut [37,38]. While we do not detect these specific strains, further, as reviewed by Lee and Kim [39], there is a growing body of evidence linking the gut microbiome to the general function of T cells in gut homeostasis. A recent study reported that CD8+ T cells can be regulated in murine guts by butyrate, a metabolite often derived from *Clostridium butyricum* [40]. Since RhCMV/SIV vaccines are known to elicit unique antigen-specific CD8+ T cell responses [1,2], these findings suggest that the differences in Clostridia communities we observed might impact or be the result of differences in RhCMV/SIV vaccine-induced T cell immunity. Closer analysis of Clostridia populations in the gut might help better understand this potential connection with RhCMV/SIV CD8+ T cell immunity and T cell immunity in general.

The identification of the vaccination group microbial signature suggests that the vaccine vectors and/or route could also impact gut microbiome compositions. We find this theory especially plausible since animals were provided with similar housing and sustenance regardless of vaccination group [31], limiting the impact of environmental and social factors on gut microbiomes. It is widely known that vaccine efficacy and immunogenicity are greatly influenced by vaccination administration strategies [41]. There are also reports that suggest associations between subclinical rhesus Cytomegalovirus (RhCMV) infection and gut microbiota [42]. While all rhesus macaques utilized in this study were naturally pre-exposed to RhCMV, there still may have been different responses to the vaccine vector that manifested in the group-associated microbiota we described. RhCMV serostatus was previously shown to affect RhCMV/SIV vaccine immunogenicity [43]; however, the vaccination regimen used differed from that of this study [31], which had a much longer pre-challenge/post-vaccination phase (91 weeks vs 36 weeks) and also used a SIV 5’ Pol expressing 68-1 RhCMV/SIV vector. At the time of writing, we are actively investigating whether serostatus impacts the efficacy of our RhCMV/SIV vaccine. In humans, CMV is estimated to be ∼83% seroprevalent [44], so improving our understanding of the relationship between RhCMV, host immunity, and the gut microbiome is critical for the clinical investigation of hCMV/HIV vaccines.

### Use of novel compositional data analysis techniques and machine learning strategies to identify gut microbial signatures

We took multiple measures to ensure the rigor of this analysis. For example, we repeatedly sampled each animal 5 times in intervals of 3 weeks to obtain reliable measurements of their gut microbiome profiles. This also allowed us to aggregate the profiles of these longitudinally collected samples from the same animal, drastically reducing data sparsity. While this is in part due to the increase in overall sequencing depth, there may have been other benefits, such as the reduction of technical zeros [45]. We evaluated major techniques of compositional data analysis, which have been shown to be better suited for 16S rRNA-based microbiome analysis in general for producing robust microbial signatures. There have been many tools developed to implement isometric log-ratios or balances [25–29], but these have very little interpretability due to their complex variance structure and have been criticized by the founder of compositional data analysis for this very reason [46]. The use of amalgamations (summed log ratios) has been introduced as a simpler alternative [47]; however, amalgamations, like balances, do not allow one to determine which individual components are changing within ratios and require expert decisions for their construction. On the other hand, CLRs are a popular standardization technique but have issues with multicollinearity in later downstream applications, such as linear modeling and correlation analysis [23,24]. We decided to develop an alternative standardization using a parsimonious reference set of ASVs and we showed that it yields the normal properties from the CLR transformation. Asymmetry is a known issue for the CLR, where co-exclusive abundance patterns lead to a skewed distribution of transformed values [22]. Adaptations to the CLR have been proposed to account for asymmetries such as the interquartile log-ratio (iqlr) and iterative iqlr [48], but these do not ensure the normal properties of CLRs nor do they address the collinearity issue. We did not observe asymmetry in our data, but it will be intriguing to assess the limitations of reference frames when asymmetries are common, e.g. in vaginal microbiome analysis where samples form community state types [49]. While it is beyond the scope of this study, a more rigorous examination of CLRs and reference frames is needed to assess how well they handle microbiome data of varying levels of diversity and sparsity.

DA analysis methods are a common approach for characterizing differences between groups, but results may drastically differ depending on the specific method chosen [17,18]. Here, we used a machine learning approach to show that we can predict RhCMV/SIV vaccine protection outcome using gut microbiome profiles and show that the microbial features we identified are corroborated by a recent compositional DA analysis method chosen because of its unique bias correction capabilities [35]. This provides additional support for the association of gut microbiomes with vaccine protection. In our analysis we benchmark CLRs, phylogenetic balances [28], and our reference standardization using random forests, a non-linear machine learning approach. We found that all types of log-ratios performed well when used as input to random forests, suggesting that in general log-ratio analysis is compatible with random forest machine learning and that our results do not depend on the specific methods used for standardization. There are recent efforts that outline strategies for using machine learning with microbiome data in general [19,20] and for precision medicine [50]. Our work here shows the potential for resolving correlates of immunity in microbiome data using these machine learning techniques. Furthermore, we show that we can dimensionally reduce microbiome data before using off-the-shelf machine learning tools by using our greedy pairwise log-ratio search, Pairbal. In this study, we used a consensus-based CV strategy, requiring ASVs to be identified in nearly all Pairbal predictive models. Such consensus-based approaches are often used to avoid overfitting and come in many different implementations, such as consensus nested CV [51]. While we observed positive results in this study, in general it would be informative to more thoroughly compare different CV approaches to assess which are most appropriate for diverse sources of 16S rRNA sequencing microbiome data.

### Limitations and future directions

There were multiple limitations of this study that provide opportunities for future investigations. For example, this study did not include baseline gut microbiome samples, so the existence of a pre-vaccination protective signature could not be ruled out. Also, the correlation we observed between the host and gut microbiome protective signatures could indicate host ‘imprinting’ in the gut microbiome, but baseline samples will be needed to verify this. Also, future work will be needed to investigate if/how the protective signature is changed when testing in rhesus macaques that are not naturally pre-exposed to RhCMV. Independent validations of this protective signature are also needed ideally using more powerful (and expensive) sequencing techniques, such as shotgun metagenomics and metatranscriptomics, which may more precisely delineate the taxonomies of these bacteria. If the protection associated microbes can be isolated, then follow-up studies using gnotobiotic models [52] and other approaches could inform the potential functions of the bacteria and possibly establish a clearer connection with CD8+ T cell immunity.

## CONCLUSIONS

In summary, to our knowledge, this study is the first to identify a pre-challenge gut microbial RhCMV/SIV protection signature, using new compositional data analysis techniques and machine learning approaches. The significant association between the gut microbiome and protection outcome we uncovered highlights the need for further investigation of the impact of gut microbiomes on RhCMV/SIV vaccination and vaccine efficacy in general.

## METHODS

### Rhesus macaques

Three vaccination groups of 15 RMs each (oral 68-1 vaccination group O, subQ 68-1 vaccination group S, subQ 68-1 + 68-1.2 group X) were utilized in this study, as previously described [31]. Rectal swab samples were collected from these animals during the pre-challenge phase (after vaccination) during weeks 76, 79, 82, 85, and 88. Samples were also collected after challenge at 28, 56, 85, 140, and 252 days post infection.

### 16S rRNA gene amplicon sequencing

DNA was extracted from cryopreserved stool and colon biopsies using the PowerFecal DNA Isolation Kit and the company’s recommended protocol (Qiagen, Valencia, CA). Sequencing libraries were prepared as described by the Earth Microbiome Project [53] and sequenced on an Illumina MiSeq Sequencer (Illumina, San Diego, CA).

### Data processing

Raw fastq files were processed using UNOISE3 implemented in USEARCH v10 [54]. For each sample, paired-end reads were merged using the fastq_mergepairs command with options optimized for our library preparation: fastq_minovlen 40, fastq_minmergelen 230, and fastq_maxmergelen 270. Reads were then filtered using a maximum expected error of 1 (fastq_maxee 1.0), which is the recommended value provided in the USEARCH manual (https://www.drive5.com/usearch/manual/exp_errs.html). PhiX was separately filtered from these reads using the filter_phix command. Next, reads were denoised using the unoise3 command to generate a set of unique ASVs. Finally, merged filtered reads were aligned and assigned to these denoised ASVs using the otutab command with the maxaccepts 10 and maxrejects 100 and top_hit_only options. Ties in alignment were broken systematically by picking the first in database file order (i.e. the more abundant ASV would be selected). ASVs without any allocated reads after this process were removed from analysis. Taxonomy of all remaining ASVs was determined using an RDP classifier with RDP database v18 [55].

The count matrix was then aggregated for each animal, separately for pre- and post-challenge samples. Samples were weighted evenly by first computing the closure of the count matrix (i.e. converting to proportions), taking the average of proportions, and by then scaling up using the total reads for the samples that were merged and rounding to produce final integer counts. This aggregated count matrix was then filtered requiring an ASV to have at least 5 counts in 50% of samples.

### Sample diversity and clustering analysis

Sample alpha diversity was computed using the Shannon diversity index. Correlations between samples were generated using Spearman correlation. The resulting Spearman correlation matrix was converted to a distance matrix by computing 1 – correlations. This was then used as input to the hclust function separately for samples from each vaccination group to perform hierarchical clustering, using the ward.D2 method. These three hierarchical clusters were then merged to form a single hierarchical cluster of all samples.

### Phylogenetic analysis

All ASVs were aligned using PRANK [56] and the resulting multiple sequence alignment was then processed in R using the phangorn package [57]. ASV sequences were clustered by pairwise sequence dissimilarity using the hclust function with the ward.D2 method and converted to a phylogenetic tree using the as.phylo function. Phylogenetic tree visualization was performed using the ggtree and dendextend R packages [58,59].

### Compositional 16S data analysis

Different types of compositional techniques were employed in this study. To handle the presence of zero counts, a pseudocount of 1 was used. Phylogenetic balances were determined using the PhILR R package [28], while centered log-ratios (CLRs) were calculated using the clr function in the compositions R package [60]. Initial principal components analysis (PCA) was performed using the top 350 most variable phylogenetic balances located at the bottom of the phylogeny (i.e. balances of size two) using the prcomp function in R. All other PCAs were also performed with the prcomp function.

Reference log-standardization was performed by first determining a reference set of ASVs, as described in the **Supplementary Methods**. ASVs were required to have a minimum count of 5 in all samples to be considered. Five iterations of the MCMC-based procedure were performed, and the optimal ASV reference set was determined by using loess regression from the R stats package with the size parameter equal to 0.5 using the variance of the ASV reference set as a function of the set size. The optimal reference set was selected as the set with a size less than or equal to the mean set size and with the most negative residual value. Pairbal was run as described in the **Supplementary Methods** using a Wilcoxon rank-sum test with a raw p-value cutoff of 0.05 and a minimum log-standardized difference of 1.

### Cross-validation using random forest machine learning

15-fold cross-validation (CV) was performed using aggregated pre-challenge samples 5 times. In each instance, random forests were generated using the randomForest R package with 5,000 trees. This was done using 3 different types of microbial features: reference log-standardized ASVs, CLRs, and phylogenetic balances. Random forest classifiers were also generated using Pairbal-selected ASVs and the most abundant ASVs, each transformed using either reference log-standardization or CLRs. The most abundant ASVs were selected as those with the highest average raw relative abundance and the number selected matched the number of Pairbal ASVs for the CV fold. For reference log-standardization, ASVs contained within the reference were excluded from this selection. Area under the curve (AUC) and receiver operator characteristic (ROC) curves were determined using the pROC R package [61]. Venn diagrams generated to compare results were produced using the VennDiagram R package [62]. A final ASV feature set was selected by requiring ASVs to be identified by Pairbal in 100% (all 75) or >95% (72 of 75) of CV predictive models for group and protection association random forest analyses, respectively.

### Differential abundance analysis of ASVs

To assess concordance between Pairbal/Random Forest-identified ASVs and those identified via differential abundance (DA) analysis, DA analysis was performed using ANCOM-BC, which treats count data as compositional [35]. The ancombc2 function, based on the CLR-transformation method, was utilized using the pre-challenge ASV count matrix as input (same as random forest analysis), disregarding structural zeros (struc_zero = FALSE) and with no specified taxonomic level (tax_level = NULL). To assess microbiome differences between the three vaccination groups and to perform similar comparisons as in the random forest analysis, a linear mixed effects model with fixed-effect term vaccination group was used thrice, comparing one group against the other two, e.g. O group versus S and X groups. To assess microbiome differences based on protection outcome, a linear mixed effects model with fixed-effect term protection was used. DA ASVs were identified from the primary result based on the selected variables for each approach. Resulting natural log-fold changes were converted to log2-fold changes (L2FCs). ASVs with L2FCs >= 1 and <= -1 were considered enriched and depleted, respectively. ASVs were found to be significant for microbiome differences between vaccination groups if their FDR adjusted p-values (Benjamini-Hochberg method) were below 0.05. Meanwhile, significant ASVs for microbiome differences associated with protection outcomes were identified when their p-values were below 0.05. For the DA analysis of vaccination groups, results from all three comparisons (O vs. S, X; S vs O, X; X vs O, S) were combined. Results from both DA analysis experiments were compared against those found via the Pairbal random forest analysis (see above).

### Permutation-based host-microbiome correlation analysis

To assess the significance of correlations between host gene expression and ASV relative abundances, a permutation-based approach was used. Correlations were computed using the biweight midcorrelation from the WGCNA R package [63]. In all cases, the primary test statistic was the median absolute value of the correlations between genes and ASVs. Three different permutations were used, each with 5,000 iterations: 1) animal label permutation, 2) random selection of ASVs, 3) random selection of genes. The first was used to assess whether the correlations between animals occurred by chance. The second assessed the significance of the particular set of ASVs chosen for correlation analysis (i.e. protection associated). The third assessed the significance of the genes used in the correlations. The genes used in this correlation analysis were Differential differentially abundant (DDE) genes, which were previously defined as genes with significantly different (absolute L2FC > 1.5 and FDR < 0.05) baseline-subtracted gene expression between protected and not protected animals [31]. Significance of test statistics were computed by determining the quantile with respect to the set of permutations, yielding empirical p-values. The configuration of ASVs and DDE genes was considered significant if all three permutation tests passed a significance threshold of 0.05.

### Statistical analysis

All statistical tests for significance for unpaired data were performed as specified (one- or two-sided), either using Wilcoxon rank-sum tests or t-tests. Pairwise comparisons were performed as specified (one- or two-sided), either using Wilcoxon signed-rank tests or paired t-tests. In analyses with multiple tests, FDR was controlled using Benjamini-Hochberg multiple hypothesis testing correction. PERMANOVA (using the adonis function from the vegan R package) was used to assess the significance of group differences in PCAs.

## DECLARATIONS

### Ethics approval and consent to participate

Rhesus macaque care and all experimental protocols and procedures were previously approved by the ONPRC Institutional Animal Care and Use Committee [31]. The ONPRC is a Category I facility. The Laboratory Animal Care and Use Program at the ONPRC is fully accredited by the American Association for Accreditation of Laboratory Animal Care and has an approved Assurance (#A3304-01) for the care and use of animals on file with the NIH Office for Protection from Research Risks. The ONPRC adheres to national guidelines established in the Animal Welfare Act (7 U.S.C. Sections 2131–2159) and the Guide for the Care and Use of Laboratory Animals (8th Edition) as mandated by the U.S. Public Health Service Policy.

### Consent for publication

Not applicable

### Availability of data and material

All 16S rRNA gene sequencing data is available under BioProject accession number PRJNA1051096 in the NCBI BioProject database (https://www.ncbi.nlm.nih.gov/bioproject/). Transcriptomic data for vaccination groups O, S, and X is available in the Gene Expression Omnibus (GEO) https://www.ncbi.nlm.nih.gov/geo/ under accession number GSE160562. **Figures S1-12** and **Table S1** are provided in **Additional File 1: Supplementary Materials**. Additional methods descriptions for the generation of microbial reference frames and the Pairbal algorithm are provided in **Additional File 2: Supplementary Methods**. Count matrices, sample metadata, ASV taxonomy, the microbial reference frame identified, and data used to generate **Figures 1c-g**, **2-6**, **S1-12** are provided in **Additional File 3: Supplementary Data**. Source code for Pairbal and a vignette showing its application are available in GitHub (https://github.com/ncsu-penglab/Pairbal).

## Authors’ contributions

This study was conceived and designed by: L.L., L.J.P., S.G.H., M.G., and X.P. NHP study and rectal swab collection were overseen by: S.G.H. 16S rRNA gene sequencing experiments were performed by: E.S. and M.C. Differential abundance analysis was performed by: S.J. and H.B. All other bioinformatics processing and analysis was performed by: H.B. The paper was written by: H.B. and X.P. The paper was reviewed and edited by all authors.

## Acknowledgements

Not applicable

