## Supplementary Materials for "Pre-challenge gut microbial signature predicts RhCMV/SIV vaccine efficacy in rhesus macaques"

H. Brochu, *et al*


| Page # | Supplementary Figure/Table |
|---|---|
| 2 | Figure S1. Hierarchical clustering of gut microbiome samples stratified by vaccination group. |
| 3 | Figure S2. Phylogenetic balance analysis of pre-challenge gut microbiomes. |
| 4 | Figure S3. Markov chain Monte Carlo (MCMC) experiments for identification of low-variance reference ASVs. |
| 5 | Figure S4. Random forest model predictions of animal vaccination groups. |
| 6 | Figure S5. Concordance between ANCOM-BC and Pairbal random forest analyses of vaccination groups. |
| 7 | Figure S6. Heatmap showing similarities of ANCOM-BC and Pairbal ASVs detected in vaccination group analyses. |
| 8 | Figure S7. Random forest model predictions of animal protection outcome. |
| 9 | Figure S8. Concordance between ANCOM-BC and Pairbal random forest analyses of animal protection outcome. |
| 10 | Figure S9. Heatmap showing similarities of ANCOM-BC and Pairbal ASVs detected in protection outcome analyses. |
| 11 | Figure S10. Heatmap showing the animal-to-animal variation of protection associated ASVs. |
| 12 | Figure S11. Phylogenetic tree of protection associated ASVs. |
| 13 | Figure S12. Permutation analysis assessing significance of correlations between differential differentially expressed (DDE) genes and protection associated ASVs. |
| 14 | Table S1. Difference in reference log-standardized values between protected and not protected animals. |

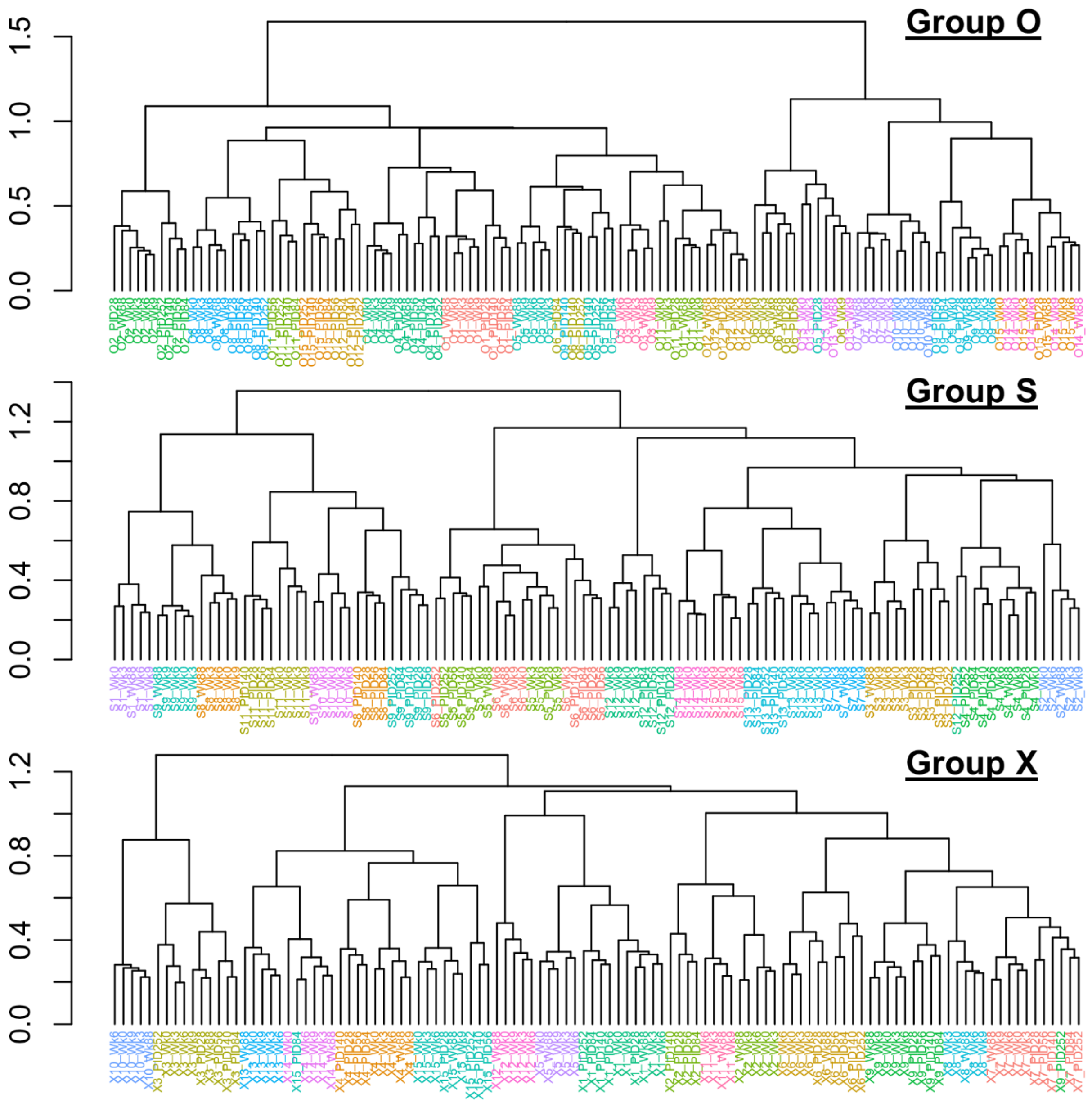


**Figure S1. Hierarchical clustering of gut microbiome samples stratified by vaccination group**. All pairwise spearman correlations were computed, and samples were clustered using a distance matrix produced from this correlation matrix. The vertical axis indicates the Euclidean distance of branches and samples are colored by their animal label.

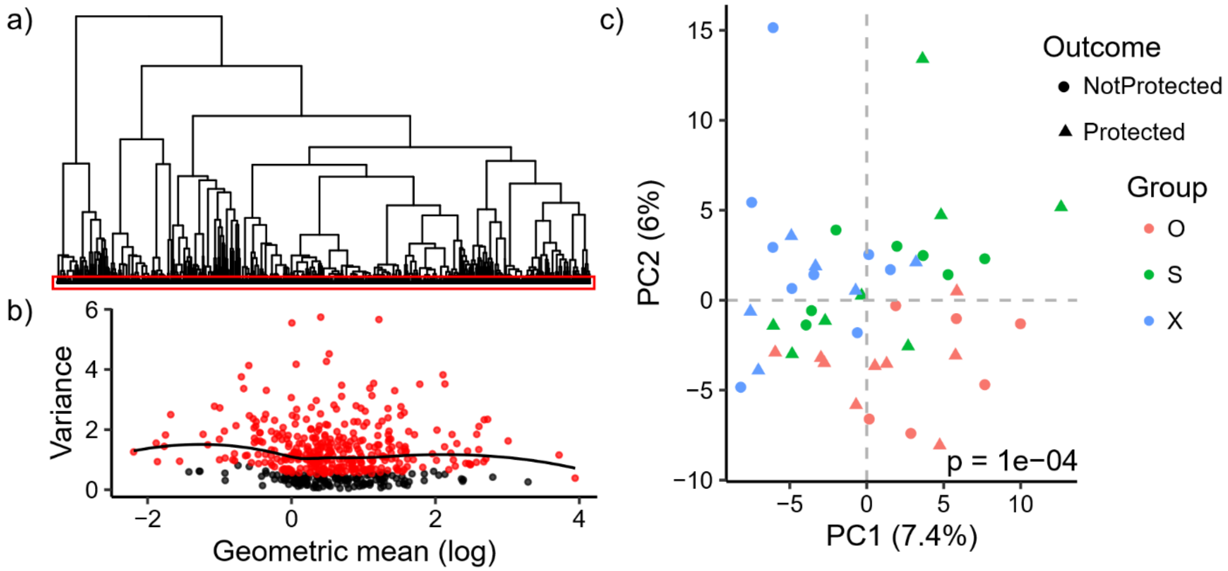


**Figure S2. Phylogenetic balance analysis of pre-challenge gut microbiomes**. **a)** Phylogenetic tree of ASVs used as orthonormal basis of phylogenetic balances with pairwise balances at the bottom of the tree indicated by a red box. **b)** Geometric mean (natural log) - variance relationship of pairwise balances with the 350 most variable balances colored red and a loess curve shown in black. **c)** Principal components analysis using pairwise balances selected in **(b)** with animals colored by group (red = O, green = S, blue = X) and shaped by challenge outcome (circle = protected, triangle = not protected). Statistical significance was determined using PERMANOVA.

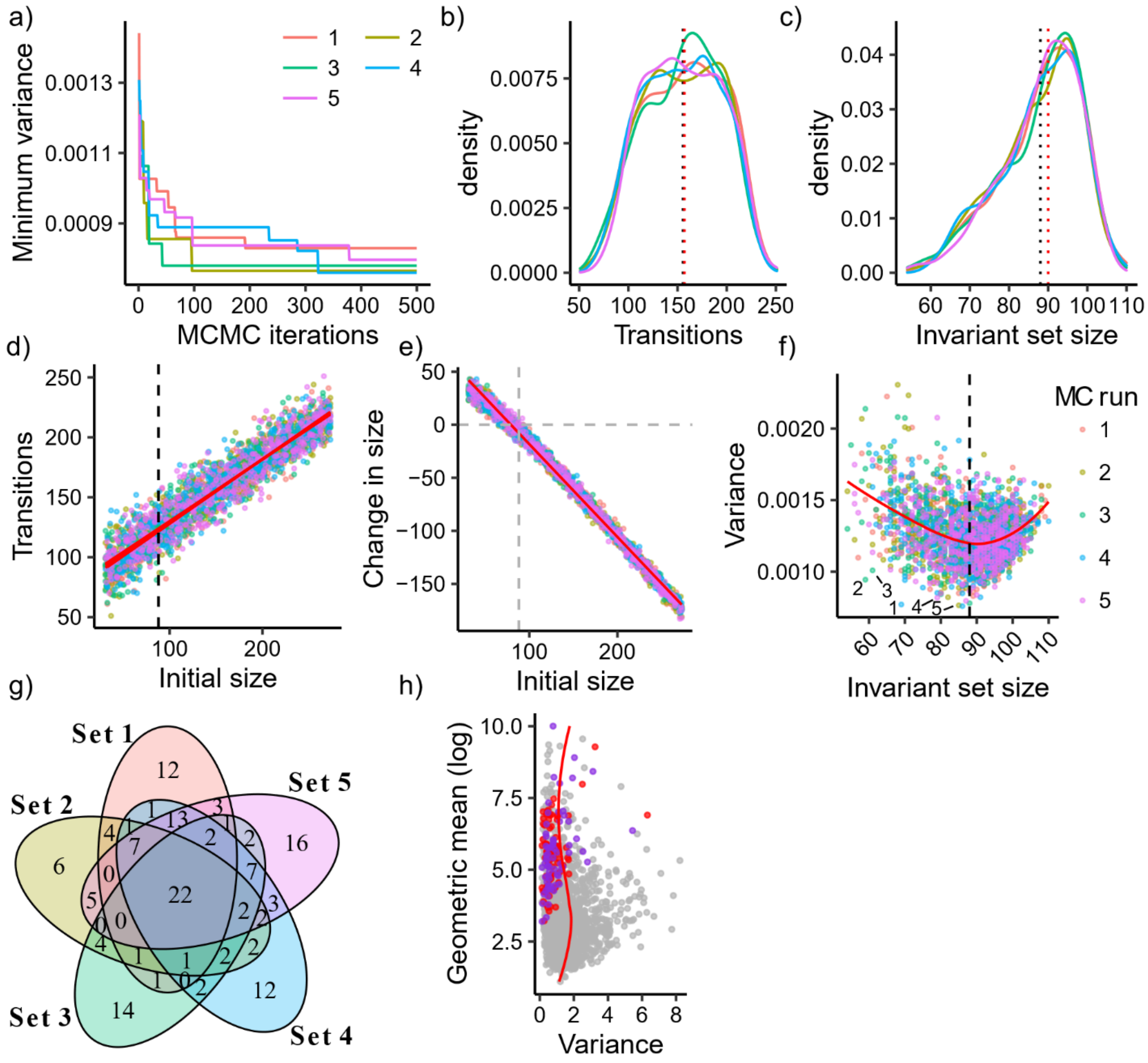


**Figure S3. Markov chain Monte Carlo (MCMC) experiments for identification of low-variance reference ASVs**. Each iteration of an MCMC experiment yields an invariant set of ASVs with a local minimum variance. In **(a-g)** the results from each experiment are colored differently. **a)** Minimum variance of all invariant sets over the course of each MCMC experiment. **b-c)** Distributions of transitions and invariant set sizes from MCMC iterations, with overall mean and median indicated by black and red dotted lines, respectively. The overall mean invariant set size is also indicated in (**d-f**) along with loess regression curves in red. **d-e)** Transitions and the change in the size of ASV sets from the initially chosen size to the final size of the invariant set. **f)** Scatter plot showing the variance and size of all invariant sets, with optimal set from each experiment (labeled 1-5) chosen using the smallest residual values. **g)** Venn diagram showing the intersections of ASVs from the optimal invariant set chosen from each MCMC experiment. **h)** ASVs are standardized using the centered log-ratio and plotted using their variance and geometric mean and a loess regression curve is shown in red. ASVs in the final invariant set, set 1 in (**g**), are shown in purple and those exclusively in the other four sets are shown in red.

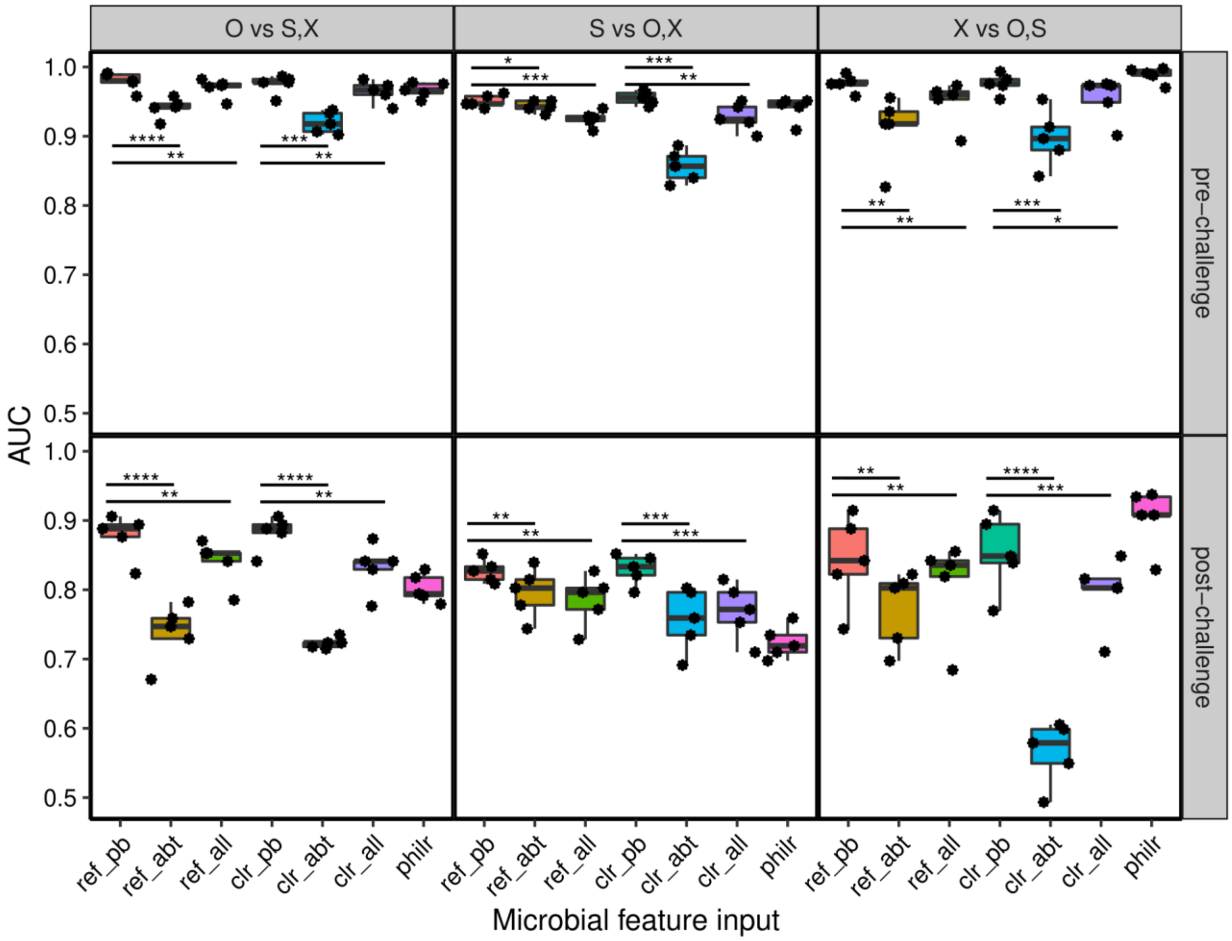


**Figure S4. Random forest model predictions of animal vaccination groups**. Area under the curve (AUC) is shown for models that predict the left out animals in the pre-challenge (top panel) and post-challenge (bottom panel) phases. Vaccination groups are grouped together for binary predictions, e.g. in O vs S,X the S and X groups are combined. Models are generated using different microbial feature inputs: phylogenetic balances (philr), reference frame standardized ASVs with all (ref_all) Pairbal (ref_pb), or most abundant (ref_abt) ASVs, or centered log-ratios with all (clr_all) Pairbal (clr_pb), or most abundant (clr_abt) ASVs. The number of most abundant ASVs used was matched to the number of ASVs identified by Pairbal. Boxplots are colored by the microbial feature input. The significance of the improvement in AUC using Pairbal was assessed using one-sided paired t-tests with FDR control. ****p<0.001, ***p<0.01, **p<0.05, *p<0.1

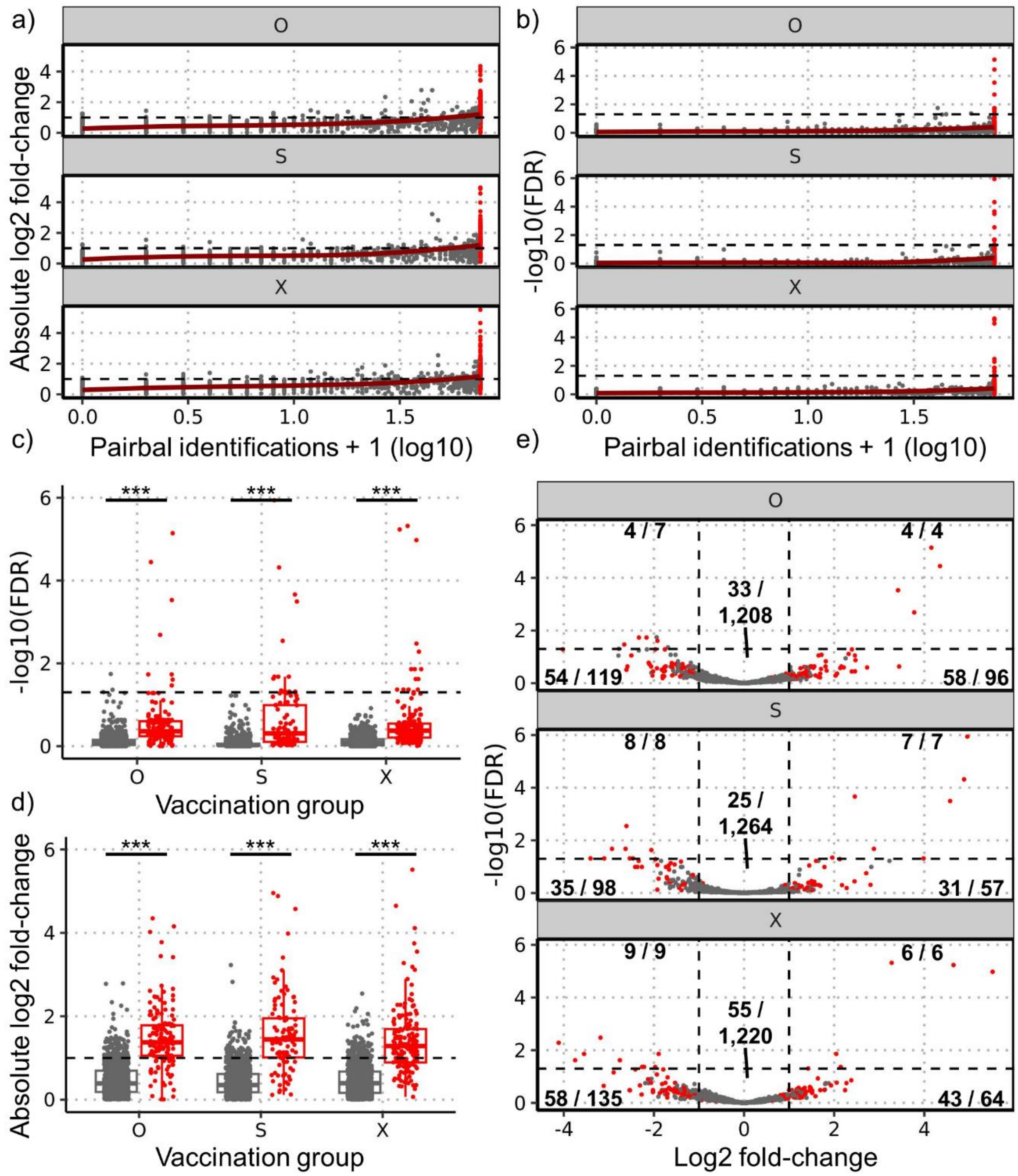


**Figure S5. Concordance between ANCOM-BC and Pairbal random forest analyses of vaccination groups**. In each analysis, vaccination groups were compared in a "one-versus-rest" fashion, e.g. "O" signifies O versus S and X. In all panels, red data points indicate ASVs detected by Pairbal in all cross-validation (CV) predictive models (consensus CV ASVs). The **(a)** absolute Log2 fold-change (L2FC) and **(b)** -log10 FDR-adjusted p-values of ASVs for each vaccination group as calculated by ANCOM-BC are plotted against Pairbal identification frequency + 1 (log10) with black horizontal dashed lines indicating the threshold for enrichment/depletion **(a)** and statistical significance **(b)**. **(c)** and **(d)** show the comparison of -log10 FDRs and absolute L2FCs, respectively, between consensus CV ASVs and all other ASVs. **e)** Volcano plot showing the L2FC and -log10 FDR of each ASV. In each region of the plot (i.e. separated by L2FC and FDR threshold dashed lines), the number of consensus CV ASVs is shown divided by the total number of ASVs. An absolute L2FC=1 is indicated by horizontal dashed lines in **b** and **d** and by vertical dashed lines indicate in **e**. A -log10 FDR=0.05 is indicated by horizontal dashed lines in **a**, **c** and **e**. Statistical significance in **c** and **d** assessed using one-sided Wilcoxon rank-sum tests with FDR control. ****p<0.001, ***p<0.01, **p<0.05, *p<0.1

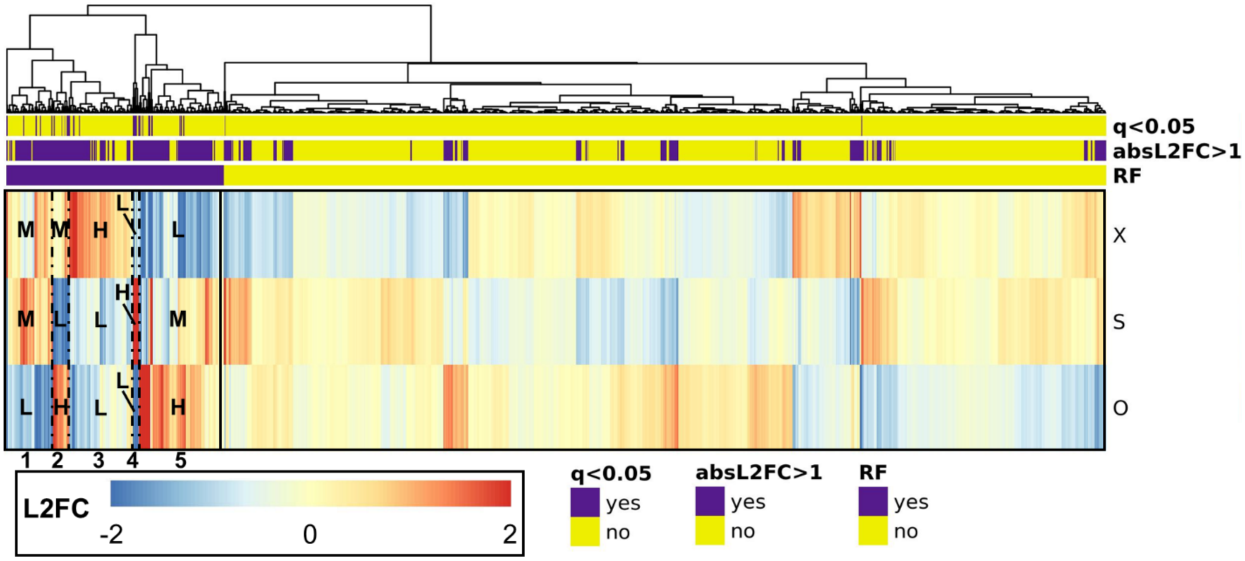


**Figure S6. Heatmap showing similarities of ANCOM-BC and Pairbal ASVs detected in vaccination group analyses**. In each analysis, vaccination groups were compared in a "one-versus-rest" fashion, e.g. "O" signifies O versus S and X. The heatmap shows log2 fold-changes (L2FCs) in each of these group comparisons fixed in the range of -2 to 2 for saturation purposes, and the ASVs are separately clustered within those selected by Pairbal and those not in any of the group comparisons (RF label above heatmap). ASVs are also labeled based on whether any of the group comparisons yielded an absolute L2FC (absL2FC) > 1 or FDR-adjusted p-value (q) < 0.05 (see **Figure S5**). ASVs selected by Pairbal are separated by a vertical solid line (left) and are further subdivided into five groups by vertical dashed lines. In each vaccination group and ASV group, a label of high (H), medium (M), or low (L) was assigned based on the relative L2FCs observed across vaccination groups.

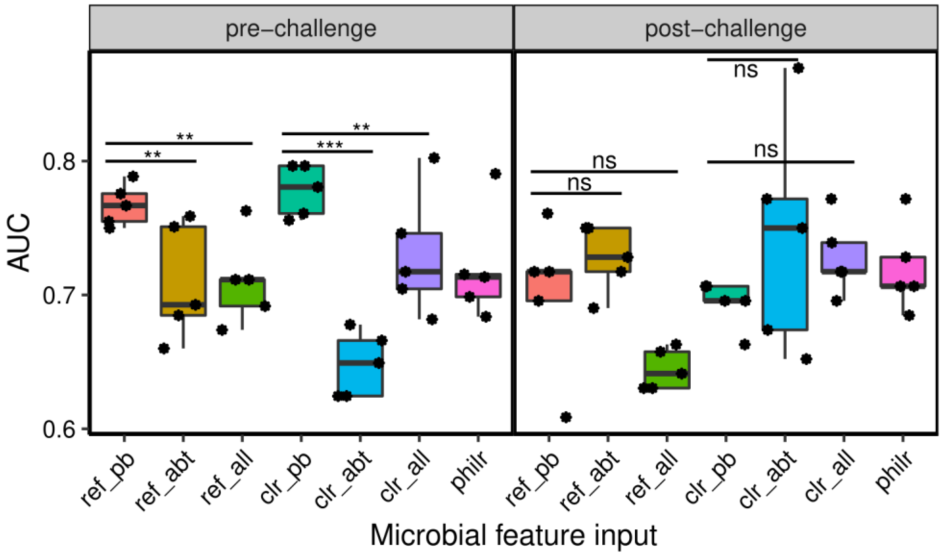


**Figure S7. Random forest model predictions of animal protection outcome**. Area under the curve (AUC) is shown for models that predict the left out animals in the pre-challenge (left panel) and post-challenge (right panel) phases. Models are generated using different microbial feature inputs: phylogenetic balances (philr), reference frame standardized ASVs with all (ref_all), Pairbal (ref_pb), or most abundant (ref_abt) ASVs, or centered log-ratios with all (clr_all), Pairbal (clr_pb), or most abundant (clr_abt) ASVs. The number of most abundant ASVs used was matched to the number of ASVs identified by Pairbal. Boxplots are colored by the microbial feature input. The significance of the improvement in AUC using Pairbal was assessed using one-sided paired t-tests with FDR control. ***p<0.01 **p<0.05, *p<0.1, ns = not significant

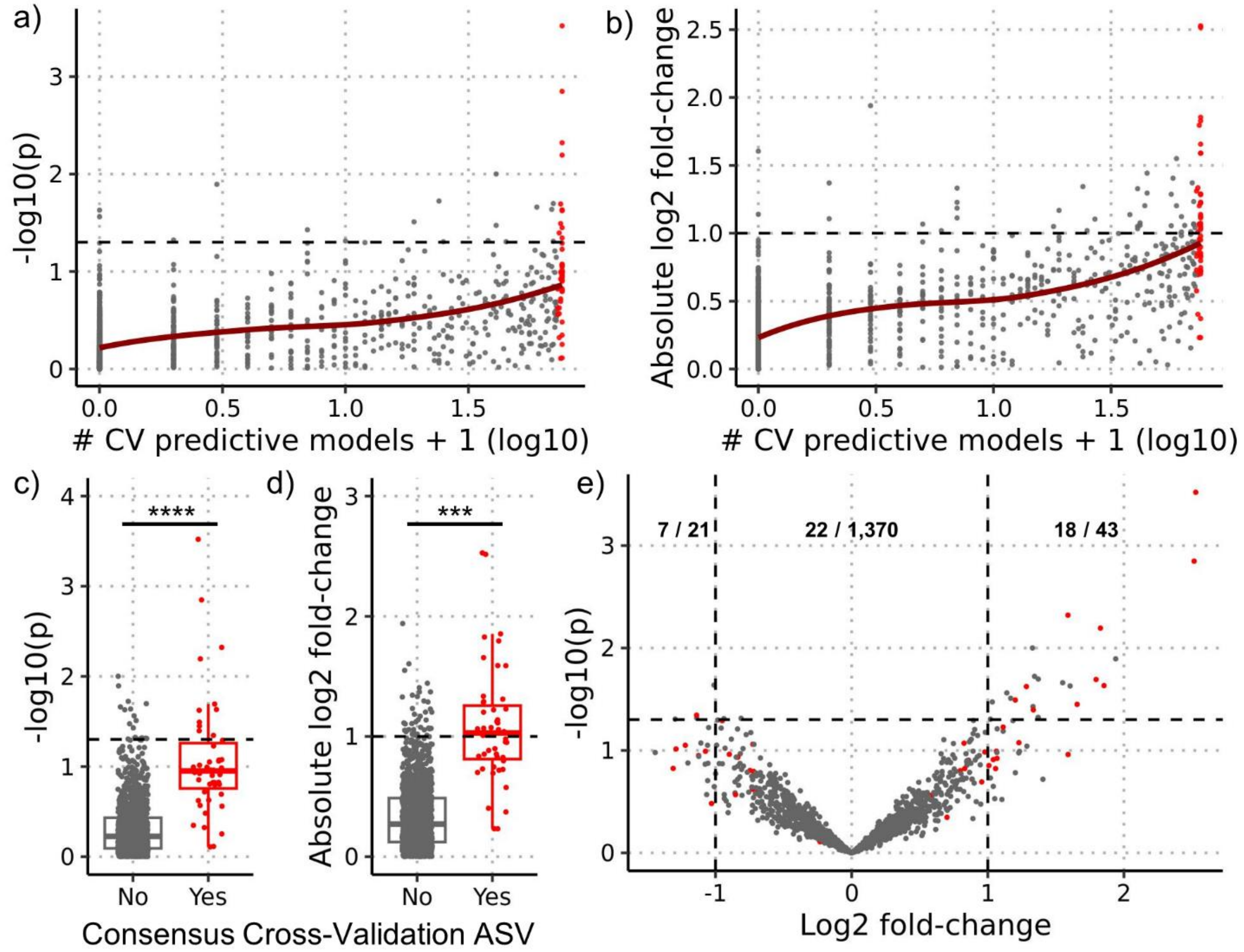


**Figure S8. Concordance between ANCOM-BC and Pairbal random forest analyses of animal protection outcome**. In each analysis, differences in ASV relative abundances were assessed by comparing protected versus no protected animals. In all panels, red data points indicate ASVs detected by Pairbal in >95% (at least 72 of 75) of cross-validation (CV) predictive models (consensus CV ASVs). The **(a)** -log10 raw p-value and **(b)** absolute log2 fold-change (L2FC) of ASVs as calculated by ANCOM-BC are plotted against the number of Pairbal identifications (log10). **(c)** and **(d)** show the comparison of p-values and absolute L2FCs, respectively, between consensus CV ASVs and all other ASVs. **e)** Volcano plot showing the L2FC and p-value of each ASV. In each region of the plot (i.e. L2FC < -1, -1 < L2FC < 1, and L2FC > 1), the number of consensus CV ASV is shown divided by the total number of ASVs. An absolute L2FC = 1 is indicated by horizontal dashed lines in **b** and **d** and by vertical dashed lines indicate in **e**. A -log10 p-value = 0.05 is indicated by horizontal dashed lines in **a**, **c** and **e**. Statistical significance in **c** and **d** was determined using one-sided Wilcoxon rank-sum tests with FDR control. ****p<0.001; ***p<0.01

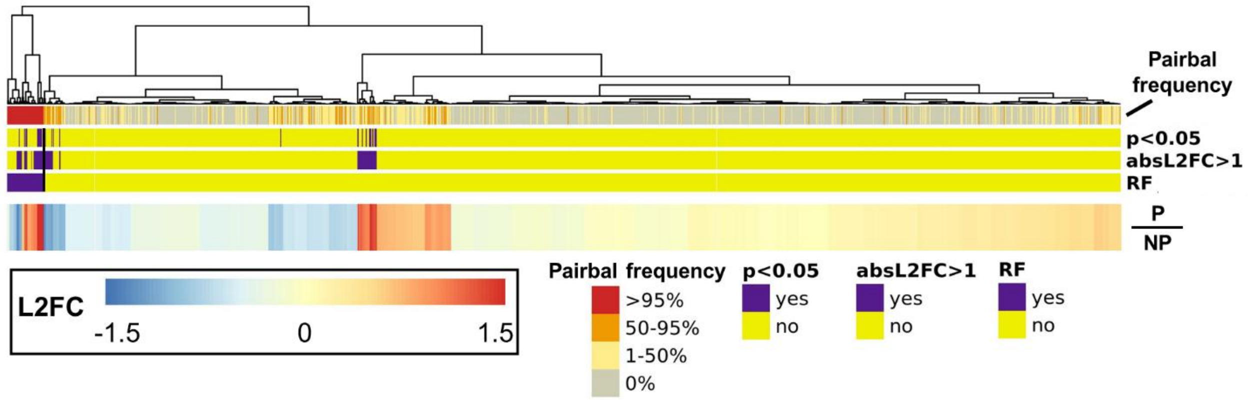


**Figure S9. Heatmap showing similarities of ANCOM-BC and Pairbal ASVs detected in protection outcome analyses**. The heatmap shows the log2 fold-changes (L2FCs) of ASVs fixed in the range of -1.5 to 1.5 for saturation purposes with ASVs clustered separately based on whether they were consensus CV ASVs or not (RF label above heatmap, see **Figure S8**). ASVs are also labeled based on whether any of the ANCOM-BC group comparisons yielded an absolute L2FC (absL2FC) > 1 or raw p-value (p) < 0.05 (**Figure S8**). ASVs are further labeled by Pairbal identification frequency (red = >95%, orange = 50-95%, yellow = 1-50%, gray = 0%). Consensus CV ASVs are separated from all other ASVs in the heatmap using a vertical solid line (left side of plot).

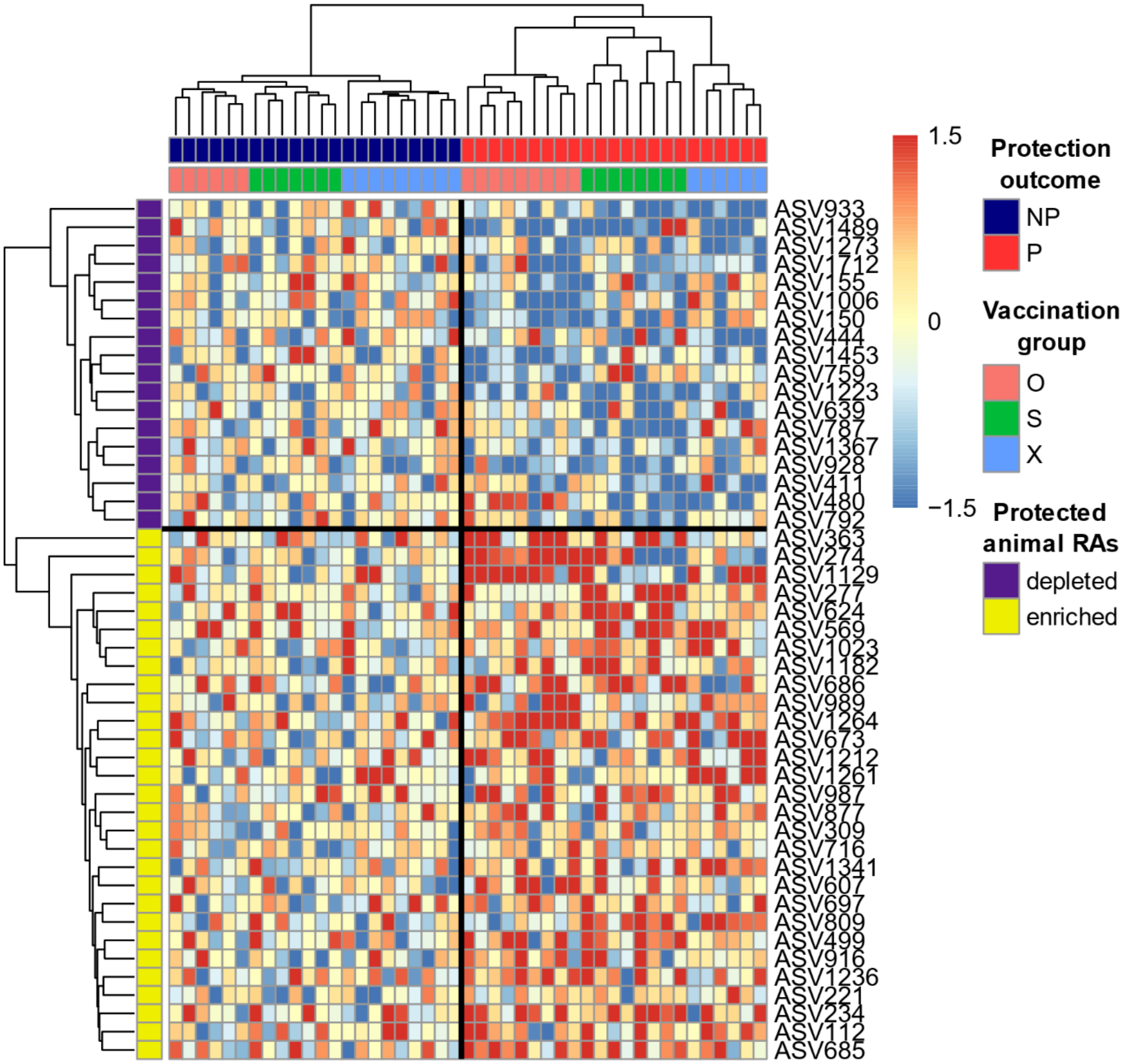


**Figure S10. Heatmap showing the pre-challenge animal-to-animal variation of protection associated ASVs**. Animals are clustered first by protection outcome (P = protected, NP = not protected), then by vaccination group. ASVs are clustered by whether they are depleted or enriched among protected animals. Log-standardized values for ASVs were scaled for each vaccination group by subtracting the median log-standardized value of NP animals and by then dividing by the standard deviation of log-standardized values for that vaccination group.

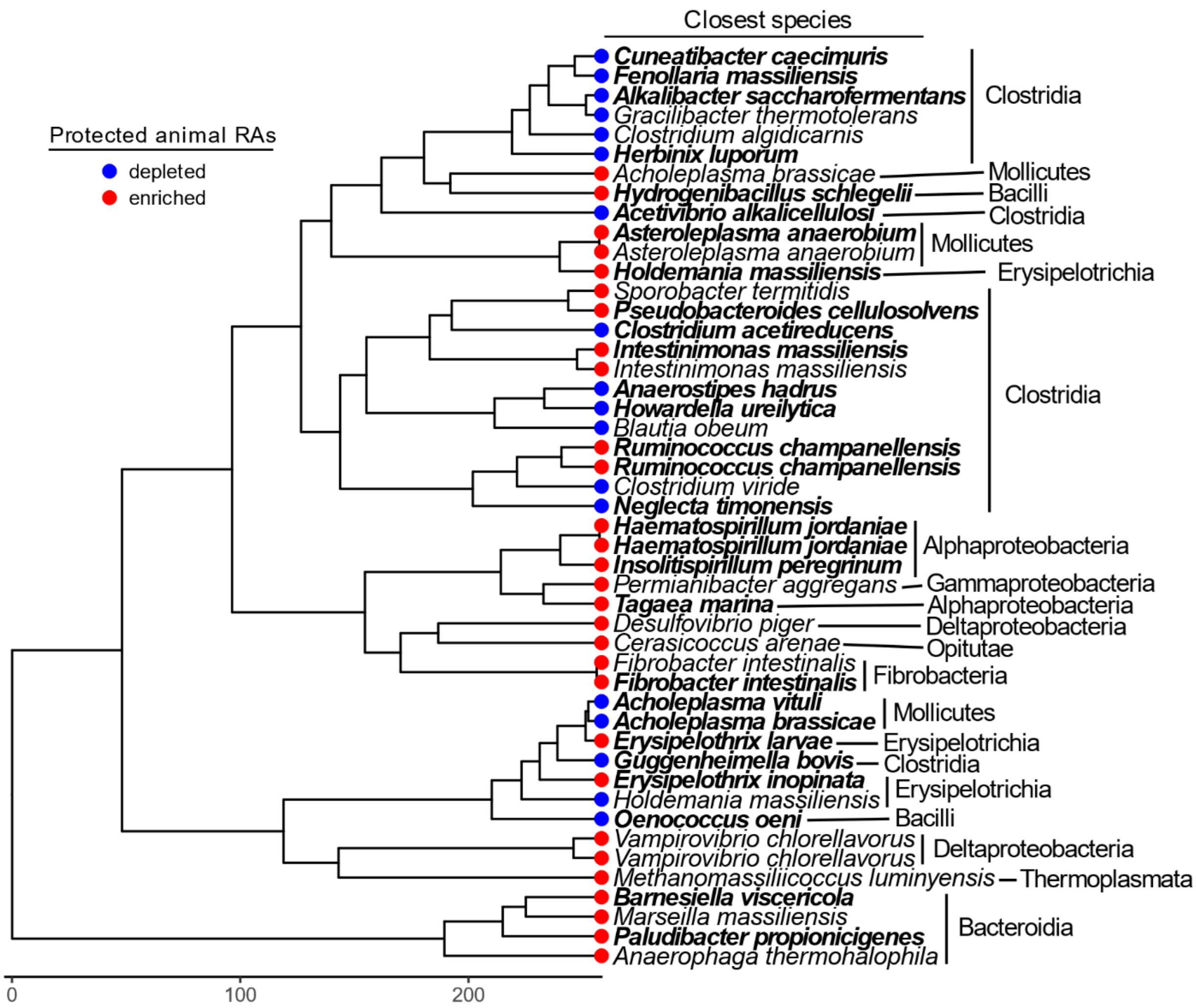


**Figure S11. Phylogenetic tree of protection associated ASVs**. Tips are colored based on whether the ASV is enriched (red) or depleted (blue) among protected animals relative to those that are not. The closest species assignment is shown to the right of the tip and further to the right the class designation is also shown. Species are in boldface if they are exclusively protection-associated (i.e. no group association); otherwise, they are in standard font.

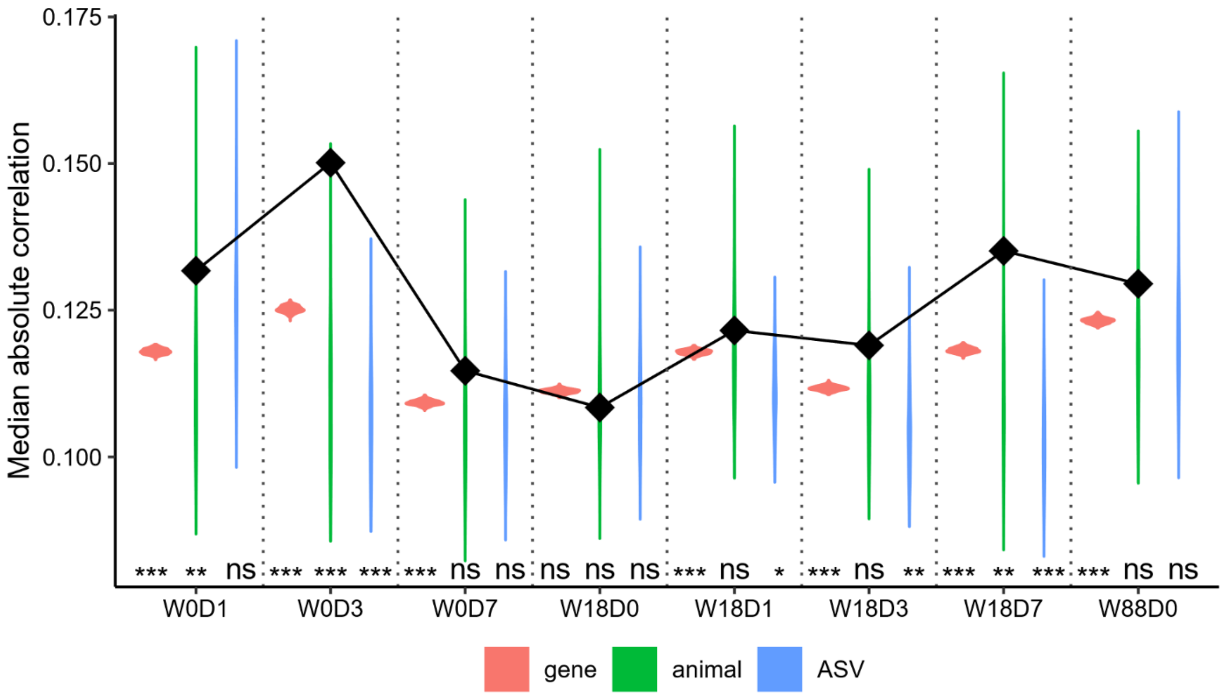


**Figure S12. Permutation analysis assessing significance of correlations between differential differentially expressed (DDE) genes and protection associated ASVs**. DDE genes are correlated with all animals and correlations are computed for all pre-challenge time points. The median absolute correlation is used as a summary statistic and corresponding null distributions for randomized animals and for randomly selected genes and ASVs are shown as violin plots. The actual computed value is signified by the black diamonds. Significance was determined by assessing the quantile of the actual summary statistic in the null distributions. *** p<0.001, ** p<0.01, * p<0.05, ns = not significant.

**Table S1. Difference in reference log-standardized values between protected and not protected animals.** Differences are shown separately for each vaccination group as well as the total average for all ASVs. Entries are colored based on the magnitude of the change (orange: > 1, yellow: between 0 and 1, light blue: between 0 and -1, dark blue: < -1). The closest species taxonomic assignment for each ASV is shown on the right.

| ASV | O | S | X | Average | Closest species |
|---|---|---|---|---|---|
| ASV234 | 1.22 | 2.35 | 0.0697 | 1.22 | *Ruminococcus champanellensis* |
| ASV569 | 0.214 | 1.81 | 1.35 | 1.12 | *Desulfovibrio piger* |
| ASV685 | 1.7 | 1.05 | 0.544 | 1.1 | *Asteroleplasma anaerobium* |
| ASV499 | 0.945 | 1.91 | 0.429 | 1.1 | *Vampirovibrio chlorellavorus* |
| ASV112 | 1.15 | 0.957 | 1.11 | 1.07 | *Paludibacter propionicigenes* |
| ASV1264 | 1.67 | 0.803 | 0.737 | 1.07 | *Ruminococcus champanellensis* |
| ASV1341 | 0.793 | 0.68 | 1.52 | 0.997 | *Cerasicoccus arenae* |
| ASV1023 | 0.787 | 1.41 | 0.671 | 0.955 | *Hydrogenibacillus schlegelii* |
| ASV916 | 0.887 | 1.22 | 0.679 | 0.93 | *Insolitispirillum peregrinum* |
| ASV673 | 1 | 1.08 | 0.642 | 0.907 | *Haematospirillum jordaniae* |
| ASV624 | 0.521 | 1.63 | 0.565 | 0.906 | *Haematospirillum jordaniae* |
| ASV1182 | 0.757 | 1.68 | 0.0891 | 0.842 | *Barnesiella viscericola* |
| ASV877 | 1.17 | 0.292 | 1.05 | 0.838 | *Tagaea marina* |
| ASV1261 | 0.37 | 0.653 | 1.32 | 0.782 | *Permianibacter aggregans* |
| ASV1236 | 1.58 | 0.633 | 0.0832 | 0.766 | *Intestinimonas massiliensis* |
| ASV221 | 0.532 | 1.3 | 0.394 | 0.742 | *Fibrobacter intestinalis* |
| ASV607 | 0.813 | 1.06 | 0.326 | 0.734 | *Holdemania massiliensis* |
| ASV989 | 0.716 | 0.362 | 0.951 | 0.676 | *Erysipelothrix larvae* |
| ASV987 | 0.695 | 0.919 | 0.229 | 0.614 | *Acholeplasma brassicae* |
| ASV363 | 2.65 | 2.07 | -0.81 | 1.3 | *Asteroleplasma anaerobium* |
| ASV697 | 1.27 | 1.79 | -0.597 | 0.822 | *Pseudobacteroides cellulosolvens* |
| ASV686 | 1.24 | 1.18 | -0.189 | 0.741 | *Intestinimonas massiliensis* |
| ASV716 | 0.532 | 1.53 | -0.13 | 0.643 | *Fibrobacter intestinalis* |
| ASV1129 | 2.24 | -0.22 | 1.15 | 1.06 | *Sporobacter termitidis* |
| ASV309 | 1.29 | -0.0298 | 1.19 | 0.818 | *Marseilla massiliensis* |
| ASV1212 | 1.5 | -0.118 | 0.523 | 0.634 | *Vampirovibrio chlorellavorus* |
| ASV274 | 2.57 | -0.316 | -0.304 | 0.649 | *Erysipelothrix inopinata* |
| ASV411 | 0.449 | -0.954 | -1.69 | -0.733 | *Guggenheimella bovis* |
| ASV480 | 1.05 | -1.89 | -1.48 | -0.774 | *Cuneatibacter caecimuris* |
| ASV277 | -0.12 | 3.88 | 1.2 | 1.65 | *Methanomassiliicoccus luminyensis* |
| ASV809 | -0.251 | 1.63 | 1.73 | 1.04 | *Anaerophaga thermohalophila* |
| ASV759 | -1.76 | 0.457 | -0.432 | -0.579 | *Holdemania massiliensis* |
| ASV1273 | -0.519 | 0.0157 | -1.6 | -0.7 | *Acholeplasma vituli* |
| ASV444 | -0.251 | 0.113 | -2.72 | -0.953 | *Anaerostipes hadrus* |
| ASV792 | -0.185 | -1.53 | 0.0709 | -0.547 | *Gracilibacter thermotolerans* |
| ASV155 | -0.837 | -1.62 | 0.354 | -0.7 | *Clostridium algidicarnis* |
| ASV1006 | -1.87 | -0.744 | 0.0302 | -0.86 | *Alkalibacter saccharofermentans* |
| ASV787 | -0.0286 | -3.01 | 0.283 | -0.919 | *Fenollaria massiliensis* |
| ASV928 | -1.42 | -1.67 | 0.312 | -0.928 | *Acholeplasma brassicae* |
| ASV150 | -2.12 | -1.91 | 0.677 | -1.12 | *Howardella ureilytica* |
| ASV1453 | -1.48 | -0.725 | -0.238 | -0.815 | *Neglecta timonensis* |
| ASV933 | -0.214 | -0.949 | -1.43 | -0.865 | *Clostridium acetireducens* |
| ASV1712 | -1.01 | -0.888 | -0.769 | -0.888 | *Oenococcus oeni* |
| ASV639 | -0.945 | -1.24 | -0.605 | -0.93 | *Clostridium viride* |
| ASV1367 | -0.892 | -1.65 | -0.248 | -0.931 | *Herbinix luporum* |
| ASV1223 | -0.521 | -1.9 | -0.837 | -1.09 | *Acetivibrio alkalicellulosi* |
| ASV1489 | -1.96 | -0.621 | -1.07 | -1.22 | *Blautia obeum* |
