## Supplementary Methods for "Pre-challenge gut microbial signature predicts RhCMV/SIV vaccine efficacy in rhesus macaques"

H. Brochu et al.

### I. Introduction

Due to the compositional nature of microbial 16S amplicon sequence data, it is becoming increasingly apparent that compositional analysis methods are needed to analyze this data [1, 2]. In short, this means that given $n$ microbial features, analysis is constrained to a simplex $\mathcal{S}^n$. Traditionally in compositional data analysis there are three types of ratios used for transforming such features: additive log-ratios (ALR), isometric log-ratios (ILR), and centered log-ratios (CLR). CLRs are a popular, interpretable transformation implemented for 16S microbiome analysis [3], but suffer from subcompositional incoherence and result in singular covariance (i.e. collinearity) [4]. These qualities make isometric log-ratios, colloquially referred to as balances, the most theoretically appropriate choice for log-ratio transformation, as it is scale invariant, permutation invariant, and subcompositionally coherent [5]. Since balances require an orthonormal basis generated by a sequential binary partition, there are many ways to implement them, resulting in numerous balance methods for microbiome analysis [6, 7, 8, 9, 10]. While an effective compositional transformation, balances have very little interpretability due to their complex variance structure. John Aitchison, the founder of compositional data analysis [4], has criticized the use of balances for this very reason [11]. A recent polemic by Michael Greenacre points out that isometric log-ratios, while mathematically beautiful constructs, have poor interpretability [12]. He further proposes the use of amalgamations (summed log ratios) as a simpler alternative [13]; however, amalgamations do not allow one to determine which individual components are changing and require expert decisions. Furthermore, balances are inherently limited since there is a loss of directionality for components when they are in ratios and one cannot determine which inidividual components are changing. The use of pairwise balances has also been shown to be an effective alternative strategy for probing compositional space [14], but to our knowledge such an approach has not yet been applied to microbial compositional data.

In this work, we propose a novel solution to identify an easily interpretable set of microbial features in compositional space standardized by a CLR-like reference frame. Herein, we describe a Markov chain Monte Carlo (MCMC)-based minimization approach to identify a low-variance set of microbial features that approximates the CLR, while also yielding the benefits of non-singular covariance. We further describe a separate strategy, called Pairbal, for enriching microbial signal of a phenotype of interest using pairwise log-ratios.

### II. Identification of reference ASVs using Markov chain Monte Carlo-based minimization

#### A. Motivation

A necessary feature of data normalization is even centering across samples, such that comparisons can be made between them. CLRs perform quite well, since by definition, they center data at 0. This can easily be shown for a sample with $n$ components of $\boldsymbol{x}$:

$$\bar{x} = \frac{1}{n}\sum_{i=0}^{n} log\frac{x_i}{g(\boldsymbol{x})} = log\frac{g(\boldsymbol{x})}{g(\boldsymbol{x})} = 0$$

In general, we use $g(\cdot)$ to indicate the geometric mean, show above as $g(\boldsymbol{x})$. While yielding desirable normalization properties, CLRs unfortunately yield collinear features, complicating downstream analysis. The ALDEx2 R package [3] offers two alternative approaches to identify low variance ASVs to be placed in the denominator. Inter-quartile log-ratios (iqlr) seek features that fall between the first and third variance quartiles in all samples. The second approach, lvha, searches for features in the bottom variance quartile in all samples. These approaches are not guaranteed to succeed (i.e. they can return too few features or none at all) and also ignore the complex covariance structure produced by the geometric mean, thus not guaranteeing the beautiful centering yielded by CLRs. Therefore, we propose a more systematic approach with the goal of producing a set of ASVs that when placed in the denominator of log-ratios can approximate the centering of CLRs.

We begin by exploring the variance of sample means, which we aim to minimize asymptotically to 0. Suppose we have a set of n ASVs, $\boldsymbol{w}$, that form a simplex $\mathcal{S}^n$. We would like to find a set of s reference ASVs, $\boldsymbol{y}$, that would form the denominators of ratios. A complementary set of ASVs from $\boldsymbol{w}$, call it $\boldsymbol{x}$, will then be the source of individual numerators, such that we form $n - s$ independent log-ratios, $z_{i/s}$. This would thus transform our data from $\mathcal{S}^n$ to $\mathbb{R}^{n-s}$.

$$z_{i/s} = log\frac{x_i}{g(\boldsymbol{y})}$$

The formulation of the mean of these ratios for each sample then resembles that of the CLR now with different components in the numerator and denominator geometric means:

$$log\frac{g(\boldsymbol{x})}{g(\boldsymbol{y})}$$

To achieve the benefits of the CLR, the above expression would need to be approximately constant. Further, this means that the *variance* of the log-ratio of the geometric means needs to be approximately 0:

$$log\frac{g(\boldsymbol{x})}{g(\boldsymbol{y})} = C \implies Var(log\frac{g(\boldsymbol{x})}{g(\boldsymbol{y})}) = 0$$

It is thus our goal to find a reference set of ASVs of size s, $\boldsymbol{y}_s$, that complements $\boldsymbol{x}_{n-s}$ such that this variance term is *minimized*:

$$\underset{\boldsymbol{y}_s}{\text{argmin}}\ Var(log\frac{g(\boldsymbol{x}_{n-s})}{g(\boldsymbol{y}_s)})$$

To address this minimization problem, we must first decompose this complex variance term. It can be done in a manner similar to that of the ILR described in [5], which is decomposed into variances of log-ratios:

$$
\begin{aligned}
Var(log\frac{g(\boldsymbol{x}_{n-s})}{g(\boldsymbol{y}_s)}) = \frac{1}{s(n-s)}\sum_{p=1}^{n-s}\sum_{q=1}^{s} var(log\frac{x_p}{y_q}) - \frac{1}{2s^2}\sum_{p=1}^{s}\sum_{q=1}^{s} var(log\frac{y_p}{y_q}) \\
- \frac{1}{2(n-s)^2}\sum_{p=1}^{n-s}\sum_{q=1}^{n-s} var(log\frac{x_p}{x_q})
\end{aligned} \tag{1}
$$

Historically, the minimization of formulae with some or many variance-like terms is challenging and often without a closed-form solution. Further, a brute force strategy also is not tractable here, as (1) quickly becomes a combinatoric nightmare as $s$ increases. These problems are in general NP-hard. With 16S compositional data we can currently expect on the order of thousands of ASVs / OTUs. One strategy to explore this complex variance landscape for minima is to use a Monte Carlo (MC)-based search, wherein we treat the variances of a given configuration of ASVs as states. To do such a search, we would require fast calculation of transitions from one to another (i.e. changes in the variance). We can imagine two possible *directions* for us to make a transition by either adding or removing from our reference set of ASVs, $\boldsymbol{y}_s$, and correspondingly removing or adding from our complementary set of ASVs, $\boldsymbol{x}_{n-s}$, to be expressed in the numerator.

### B. Derivation of log-ratio variance transitions

Before we begin we must first compute the log-ratio covariance matrix, requiring $\mathcal{O}(p \cdot n^2)$ computations, where $p$ is our sample size and $n$ is the number of ASVs. This matrix can then efficiently be queried during our MC procedure. Here, we show the example of adding an ASV to $\boldsymbol{y}_s$ and later show the corresponding solution to substraction of an ASV. In either case, we begin with a proposed set of ASVs of size $s$ at iteration $i = 0$, $\boldsymbol{y}_{s_0}$, with variance denoted $V_0$. By adding ASV z to this set, we arrive at a new variance state, $V_1^{+z}$. We must now identify the change in variance. Using (1), we can show that the variance of the initial state:

$$
\begin{aligned}
V_0 = \frac{1}{(s_0)(n-s_0)}\sum_{p=1}^{n-s_0}\sum_{q=1}^{s_0} var(log\frac{x_p}{y_q}) - \frac{1}{2s_0^2}\sum_{p=1}^{s_0}\sum_{q=1}^{s_0} var(log\frac{y_p}{y_q}) \\
- \frac{1}{2(n-s_0)^2}\sum_{p=1}^{n-s_0}\sum_{q=1}^{n-s_0} var(log\frac{x_p}{x_q})
\end{aligned} \tag{2}
$$

Using (1), we can also show the variance of the proposed new state in terms of $\boldsymbol{y}_{s_0}$, $\boldsymbol{x}_{n-s_0}$, and $z$:

$$
\begin{aligned}
V_1^{+z} = &\frac{1}{(s_0+1)(n-s_0-1)}\Big[\sum_{p=1}^{n-s_0}\sum_{q=1}^{s_0} var(log\frac{x_p}{y_q}) + \sum_{p=1}^{n-s_0} var(log\frac{z}{x_p}) - \sum_{p=1}^{s_0} var(log\frac{z}{y_p})\Big] \\
&- \frac{1}{2(s_0+1)^2}\Big[\sum_{p=1}^{s_0}\sum_{q=1}^{s_0} var(log\frac{y_p}{y_q}) + 2\sum_{p=1}^{s_0} var(log\frac{z}{y_p})\Big] \\
&- \frac{1}{2(n-s_0-1)^2}\Big[\sum_{p=1}^{n-s_0}\sum_{q=1}^{n-s_0} var(log\frac{x_p}{x_q}) - 2\sum_{p=1}^{n-s_0} var(log\frac{z}{x_p})\Big]
\end{aligned}
\tag{3}
$$

Note that we do not need to worry about counting variances of log-ratios between $z$ and $x_p$, where $x_p = z$, since such terms are equal to zero. To calculate the change in variance, we can subtract (2) from (3) and combine variance terms:

$$
\begin{aligned}
V_1^{+z} - V_0 = &\Big[\frac{1}{(s_0+1)(n-s_0-1)} - \frac{1}{s_0(n-s_0)}\Big]\sum_{p=1}^{n-s_0}\sum_{q=1}^{s_0} var(log\frac{x_p}{y_q}) \\
&+ \Big[\frac{1}{2s_0^2} - \frac{1}{2(s_0+1)^2}\Big]\sum_{p=1}^{s_0}\sum_{q=1}^{s_0} var(log\frac{y_p}{y_q}) \\
&+ \Big[\frac{1}{2(n-s_0)^2} - \frac{1}{2(n-s_0-1)^2}\Big]\sum_{p=1}^{n-s_0}\sum_{q=1}^{n-s_0} var(log\frac{x_p}{x_q}) \\
&- \frac{n}{(s_0+1)^2(n-s_0-1)}\sum_{p=1}^{s_0} var(log\frac{z}{y_p}) + \frac{n}{(s_0+1)(n-s_0-1)^2}\sum_{p=1}^{n-s_0} var(log\frac{z}{x_p})
\end{aligned}
\tag{4}
$$

We see that (4) has constant coefficents that are functions of $n$ and the current state size, $s_0$. Further, the first three variance terms can be computed from the current state, while the remaining two are vectors with lengths of $\boldsymbol{z}$. The number of possible ASVs we can add is limited by the length of our current numerator set, $\boldsymbol{x}_{n-s}$. In 16S microbiome datasets, it might be beneficial to restrict the number of candidates for inclusion in the reference set of ASVs based on abundance or the presence of zero counts across samples. If we choose $k$ candidate ASVs for our search such that $k \leq n$, we can write a generalized form of (4) for the transition resulting in z being added to $\boldsymbol{y}_s$:

$$
\begin{aligned}
&\boldsymbol{t}^{+z}_{i,k-s_i} = V^{+z}_{i+1} - V_i = \alpha^+_i A_i + \beta^+_i B_i + \gamma^+_i C_i + \delta^+_i \boldsymbol{D}^+_{i,k-s_i} + \varepsilon^+_i \boldsymbol{E}^+_{i,k-s_i} \\
&\alpha^+_i = \frac{1}{(s_i+1)(n-s_i-1)} - \frac{1}{s_i(n-s_i)} \,,\ \beta^+_i = \frac{1}{2s_i^2} - \frac{1}{2(s_i+1)^2} \,, \\
&\gamma^+_i = \frac{1}{2(n-s_i)^2} - \frac{1}{2(n-s_i-1)^2} \,,\ \delta^+_i = \frac{-n}{(s_i+1)^2(n-s_i-1)} \,, \\
&\varepsilon^+_i = \frac{n}{(s_i+1)(n-s_i-1)^2} \,, \\
&A_i = \sum_{p=1}^{n-s_i}\sum_{q=1}^{s_i} var(log\frac{x_p}{y_q}) \,,\ B_i = \sum_{p=1}^{s_i}\sum_{q=1}^{s_i} var(log\frac{y_p}{y_q}) \,,\ C_i = \sum_{p=1}^{n-s_i}\sum_{q=1}^{n-s_i} var(log\frac{x_p}{x_q}) \,, \\
&\boldsymbol{D}^+_{i,k-s_i} = \sum_{p=1}^{s_i} var(log\frac{\boldsymbol{z}_{k-s_i}}{y_p}) \,,\ \boldsymbol{E}^+_{i,k-s_i} = \sum_{p=1}^{n-s_i} var(log\frac{\boldsymbol{z}_{k-s_i}}{x_p})
\end{aligned} \tag{5}
$$

Similarly, we can adapt (5) for the *removal* of ASV $z$ from $\boldsymbol{y}_s$:

$$
\begin{aligned}
&\boldsymbol{t}^{-z}_{i,s_i} = V^{-z}_{i+1} - V_i = \alpha^-_i A_i + \beta^-_i B_i + \gamma^-_i C_i + \delta^-_i \boldsymbol{D}^-_{i,s_i} + \varepsilon^-_i \boldsymbol{E}^-_{i,s_i} \\
&\alpha^-_i = \frac{1}{(s_i-1)(n-s_i+1)} - \frac{1}{s_i(n-s_i)} \,,\ \beta^-_i = \frac{1}{2s_i^2} - \frac{1}{2(s_i-1)^2} \,, \\
&\gamma^-_i = \frac{1}{2(n-s_i)^2} - \frac{1}{2(n-s_i+1)^2} \,,\ \delta^-_i = \frac{n}{(s_i-1)^2(n-s_i+1)} \,, \\
&\varepsilon^-_i = \frac{-n}{(s_i-1)(n-s_i+1)^2} \,, \\
&A_i = \sum_{p=1}^{n-s_i}\sum_{q=1}^{s_i} var(log\frac{x_p}{y_q}) \,,\ B_i = \sum_{p=1}^{s_i}\sum_{q=1}^{s_i} var(log\frac{y_p}{y_q}) \,,\ C_i = \sum_{p=1}^{n-s_i}\sum_{q=1}^{n-s_i} var(log\frac{x_p}{x_q}) \,, \\
&\boldsymbol{D}^-_{i,s_i} = \sum_{p=1}^{s_i} var(log\frac{\boldsymbol{z}_{s_i}}{y_p}) \,,\ \boldsymbol{E}^-_{i,s_i} = \sum_{p=1}^{n-s_i} var(log\frac{\boldsymbol{z}_{s_i}}{x_p})
\end{aligned} \tag{6}
$$

The terms in (5) and (6) are dependent on the removal/addition of an ASV, with the exception of $A_i$, $B_i$ and $C_i$, which are computed based on the current state.

### C. Monte Carlo approach for log-ratio variance minimization

Monte Carlo sampling strategies are common for iterative random sampling. In the previous section, we delineated variance transitions in the log-ratio variance landscape, resulting in (5) and (6). Here, we will adapt these equations to form proposal probability distributions for markov chains, enabling semi-random traversal through log-ratio variances. Each iteration of this approach will result in stopping the chain at a local minimum. A prerequisite for this algorithm is the $nxn$ log-ratio covariance matrix, $\boldsymbol{X}_{nxn}$, which can easily be computed prior to initiating this approach.

We begin an iteration by randomly selecting a set of ASVs of size $s_0$, $\boldsymbol{y}_{s_0}$, from our pool of $k$ candidate ASVs, where $k \leq n$. Using (2), we compute the variance, $V_0$, of this ASV configuration and also record the values of $A_0$, $B_0$, and $C_0$ which form the decomposition of the variance and are described in (5) and (6). Finally, we must also generate the possible transitions from scratch by computing the remaining terms in (5) and (6). In total, this setup requires $\mathcal{O}(n^2)$ lookups in our log-ratio covariance matrix.

Using the terms in (5) and (6), we can compute the possible steps in variance by the addition or removal of an ASV to our reference set, $\boldsymbol{y}_{s_0}$, respectively denoted $\boldsymbol{t}^{+z}_{0,k-s_0}$ and $\boldsymbol{t}^{-z}_{0,s_0}$. Note that in the edge cases where $s = k$ or $s = 1$, we will only consider removal and addition of an ASV, respectively. From the possible steps in variance, we only consider those that would result in a reduction in variance, i.e. only negative entries. By concatenating the two vectors containing possible steps, we can generate a single adjusted vector, $\widetilde{\boldsymbol{t}}_0$, of length $k$:

$$\widetilde{\boldsymbol{t}}_{0,j} = \begin{cases} \boldsymbol{t}_{0,j}, & \text{if } \boldsymbol{t}_{0,j} < 0 \\ 0, & \text{otherwise} \end{cases} \tag{7}$$

If $\sum \widetilde{\boldsymbol{t}}_0 = 0$, then there are no possible reductions in variance and we have reached a local minimum. At this point, we stop the iteration and record the current reference set of ASVs, $\boldsymbol{y}_{s_0}$. If at least one term is negative, then we proceed by generating a probability for each possible step proportional to their magnitude:

$$\widetilde{\boldsymbol{p}}_{0,j} = \frac{\widetilde{\boldsymbol{t}}_{0,j}}{\sum_{j=1}^{k} \widetilde{\boldsymbol{t}}_{0,j}} \tag{8}$$

Finally, we make our step by randomly drawing from the probability distribution defined by (8), corresponding to the addition or removal of ASV $z^*$ to/from our reference set.

The beauty of this algorithm lies in the subsequent steps in this markov chain, where swift updates can be made to the variance terms. We will first consider how these updates are made in general after the *addition* of an ASV $z^*$ to our reference set $\boldsymbol{y}_{s_i}$, resulting in $\boldsymbol{y}_{s_{i+1}}$, where $s_{i+1} = s_i + 1$:

$$\begin{aligned}
&\boldsymbol{z}_{s_{i+1}} = \langle \boldsymbol{z}_{s_i},\ z^* \rangle,\ \boldsymbol{z}_{k-s_{i+1}} = \boldsymbol{z}_{k-s_i}(\boldsymbol{z}_{k-s_i} \neq z^*) \\
&A_{i+1} = A_i - \boldsymbol{D}^{+}_{i,k-s_i}(z^*) + \boldsymbol{E}^{+}_{i,k-s_i}(z^*) \\
&B_{i+1} = B_i + 2\boldsymbol{D}^{+}_{i,k-s_i}(z^*) \\
&C_{i+1} = C_i - 2\boldsymbol{E}^{+}_{i,k-s_i}(z^*) \\
&\boldsymbol{D}^{+}_{i,k-s_{i+1}} = \boldsymbol{D}^{+}_{i,k-s_i}(\boldsymbol{z}_{k-s_{i+1}}) + var(log\frac{\boldsymbol{z}_{k-s_{i+1}}}{z^*}) \\
&\boldsymbol{E}^{+}_{i,k-s_{i+1}} = \boldsymbol{E}^{+}_{i,k-s_i}(\boldsymbol{z}_{k-s_{i+1}}) - var(log\frac{\boldsymbol{z}_{k-s_{i+1}}}{z^*}) \\
&\boldsymbol{D}^{-}_{i,s_{i+1}} = \langle \boldsymbol{D}^{-}_{i,s_i},\ \sum_{p=1}^{s_i} var(log\frac{z^*}{y_p}) \rangle + var(log\frac{\boldsymbol{z}_{s_{i+1}}}{z^*}) \\
&\boldsymbol{E}^{-}_{i,s_{i+1}} = \langle \boldsymbol{E}^{-}_{i,s_i},\ \sum_{p=1}^{n-s_i} var(log\frac{z^*}{x_p}) \rangle - var(log\frac{\boldsymbol{z}_{s_{i+1}}}{z^*})
\end{aligned} \tag{9}$$

Notably, all of the terms in (9) can collectively be computed in $\mathcal{O}(n)$. To compute $\boldsymbol{t}^{+}_{i+1,s_{i+1}}$ and $\boldsymbol{t}^{-}_{i+1,s_{i+1}}$ the coefficients for variance terms also need to be updated using our new reference size, $s_{i+1}$, which can be done in constant time. Below, we show the updated variance terms if an ASV $z^*$ is *removed* from the reference set $\boldsymbol{y}_{s_i}$, resulting in $\boldsymbol{y}_{s_{i+1}}$, where $s_{i+1} = s_i - 1$:

$$\begin{aligned}
&\boldsymbol{z}_{s_{i+1}} = \boldsymbol{z}_{s_i}(\boldsymbol{z}_{s_i} \neq z^*),\ \boldsymbol{z}_{k-s_{i+1}} = \langle \boldsymbol{z}_{k-s_i},\ z^* \rangle \\
&A_{i+1} = A_i + \boldsymbol{D}^{-}_{i,k-s_i}(z^*) - \boldsymbol{E}^{-}_{i,k-s_i}(z^*) \\
&B_{i+1} = B_i - 2\boldsymbol{D}^{-}_{i,k-s_i}(z^*) \\
&C_{i+1} = C_i + 2\boldsymbol{E}^{-}_{i,k-s_i}(z^*) \\
&\boldsymbol{D}^{+}_{i,k-s_{i+1}} = \langle \boldsymbol{D}^{+}_{i,k-s_i}, \sum_{p=1}^{s_i} var(log\frac{z^*}{y_p}) \rangle - var(log\frac{\boldsymbol{z}_{k-s_{i+1}}}{z^*}) \\
&\boldsymbol{E}^{+}_{i,k-s_{i+1}} = \langle \boldsymbol{E}^{+}_{i,k-s_i}, \sum_{p=1}^{n-s_i} var(log\frac{z^*}{x_p}) \rangle + var(log\frac{\boldsymbol{z}_{k-s_{i+1}}}{z^*}) \\
&\boldsymbol{D}^{-}_{i,s_{i+1}} = \boldsymbol{D}^{-}_{i,s_i}(\boldsymbol{z}_{s_{i+1}}) - var(log\frac{\boldsymbol{z}_{s_{i+1}}}{z^*}) \\
&\boldsymbol{E}^{-}_{i,s_{i+1}} = \boldsymbol{E}^{-}_{i,s_i}(\boldsymbol{z}_{s_{i+1}}) + var(log\frac{\boldsymbol{z}_{s_{i+1}}}{z^*})
\end{aligned} \tag{10}$$

In many ways the terms in (10) reflect those in (9), where we see signs flipped. This transition also requires $\mathcal{O}(n)$ computations to update terms. Once the terms are accordingly computed based on the transition, we can proceed with adjusting the steps as in (7). As mentioned before, if $\sum \widetilde{\boldsymbol{t}}_i = 0$, then this iteration is halted and the results are saved, as we have reached a local minimum. If not, then step probabilities are computed using (8), a step is drawn from this distribution, and we once again update based on the type of step (i.e. addition or removal of an ASV from the reference set of ASVs). This is repeated until a local minimum is reached.

Depending on the number of ASVs, $k$, selected as candidates for the reference set, many MC iterations may be needed to adequately sample and identify local minima. Based on the $\mathcal{O}(n^2)$ setup computations and the $\mathcal{O}(n)$ transition computations, we can expect an entire iteration to take $\mathcal{O}(max(n^2\,,\ c \cdot n))$, where $c$ is the average number of iterations to reach a local minimum. The value of $c$ could vary significantly depending on the number of candidate ASVs and might be larger than $n$ in some cases. If we run $m$ MC iterations, the final asymptotic complexity for this algorithm is in $\mathcal{O}(m \cdot max(n^2, c \cdot n))$. Note that before running the MC procedure, we required $\mathcal{O}(p \cdot n^2)$ computations for the log-ratio covariance matrix, where $p$ is our sample size. In general, we can expect $p < m$ for typical microbiome studies, thus yielding a final complexity matching that of the MC procedure: $\mathcal{O}(m \cdot max(n^2, c \cdot n))$. While this complexity does not directly depend on the number of candidate ASVs, $k$, one might expect a longer convergence time, $c$, to reach local minima or a greater number of MC iterations, $m$, to adequately sample the variance landscape when $k$ is increased.

### III. Greedy microbial feature enrichment using Pairbal

Feature selection is a challenging problem when the number of features greatly outnumber the samples available for analysis. This is a common issue for 16S microbiome data and biological data in general, with the exception of single-cell sequencing. We aim to address this problem here by probing compositional space with pairwise log-ratios without the need for a reference frame. This reference-free approach mitigates bias to any asymmetries in the simplex and allows one to apply any standardization to enriched features following the procedure, although we encourage the use of MCMC-based reference frames described above.

We begin by computing all pairwise log-ratios of features in the dataset, requiring $\mathcal{O}(n^2)$ computations. If count data is sparse, we encourage rigorous pre-filtering to reduce the computations needed at this step. We then filter these based on how well they segregate samples along our phenotype of interest. In this study, we consider the simple case of a strictly binary phenotype and generate raw p-values (wilcoxon rank-sum test) and mean differences for all pairwise log-ratios. Pairwise log-ratios are then filtered with user-defined cutoffs for these criteria.

Next, we assign these filtered pairwise log-ratios to each feature in all pairs. Thus, each feature obtains a count associated with its frequency among these pairwise log-ratios, yielding a total frequency of exactly twice the number of pairwise log-ratios. Our next and final step is to *prune* the list of features with non-zero pairwise log-ratio counts, such that each log-ratio is only assigned to a single feature. This step is motivated by the rationale that pairwise log-ratios might be more strongly influenced by only one of the two constituent features. It is here that we propose a greedy approach, where pairwise log-ratios are only counted for the feature with the higher frequency. This is done progressively, beginning with pairwise log-ratios assigned to the most frequently occurring feature. Features with non-zero counts are then kept for downstream analysis, e.g. secondary feature selection and/or modeling using machine learning applications.

### IV. Concluding remarks

Compositional data analysis has spurred a variety of approaches for representing data in compositional space, circling the central tenet of log-ratio-based analysis. Many new approaches avoid the CLR in part due to the issue of collinearity, and instead identify complex ratios known as balances. These ratios offer superior mathematical properties, but require a sequential binary partition (i.e. binary tree) for construction. The PhILR [6] and Phylofactor [7] methods use perhaps the most practical choice by generating balances from phylogenies; however, the resulting balances are challenging to interpret. We describe a simple alternative that achieves CLR-like centering while avoiding the symmetric interdependence of CLRs. This is achieved through generation of a reference set of ASVs that is used exclusively for standardization of all other ASVs. We regard this as a superior approach due to the ease of interpretation and representation of ASVs individually in compositional space with respect to the reference frame. Like balances, this transformation into real space enables safe application of standard statistical techniques and feature selection routines. We encourage the use of Pairbal in combination with other feature selection routines, especially as the number of microbial features may greatly outnumber the samples used.

Increasing the resolution of 16S rRNA amplicon sequencing by replacing OTUs with ASVs is inexorable. Beyond the improvements in reproducibility, this heightened resolution greatly improves our chances of finding microbial features common to all samples. With future improvements in sequencing depth, we anticipate that this approach will become an increasingly viable alternative to other standard compositional transformation techniques. Further research is needed to assess whether this approach is appropriate for highly asymmetric and/or less diverse microbiomes from different body sites.
